# Cortical cerebrovascular and metabolic perturbations in the 5xFAD mouse model of Alzheimer’s disease

**DOI:** 10.1101/2023.05.05.539629

**Authors:** Amandine Jullienne, Jenny I. Szu, Ryan Quan, Michelle V. Trinh, Tannoz Norouzi, Brenda P. Noarbe, Amanda A. Bedwell, Kierra Eldridge, Scott C. Persohn, Paul R. Territo, Andre Obenaus

**Affiliations:** Department of Pediatrics, School of Medicine, University of California, Irvine, Irvine, CA, USA; Department of Medicine, School of Medicine, Indiana University, Indianapolis, IN, USA; Stark Neuroscience Research Institute, School of Medicine, Indiana University, Indianapolis, IN, USA

**Author notes:** Correspondence: Andre Obenaus, PhD.

**Keywords:** Cerebrovasculature, PET, MRI, Sex, Blood-brain barrier, vessels

## Abstract

The 5xFAD mouse model is a popular model of familial Alzheimer’s Disease (AD) that is characterized by early beta-amyloid (Aβ) deposition and cognitive decrements. Despite numerous studies, the 5xFAD mouse has not been comprehensively phenotyped for vascular and metabolic perturbations over its lifespan. Male and female 5xFAD and WT littermates underwent *in vivo* ^18^F-Fluorodeoxyglucose (FDG) positron emission tomography (PET) imaging at 4, 6, and 12 months of age to assess regional glucose metabolism. A separate cohort of mice (4, 8, 12 months) underwent “vessel painting” which labels all cerebral vessels and were analyzed for vascular characteristics such as vessel density, junction density, vessel length, network complexity, number of collaterals and vessel diameter. With increasing age, vessels on the cortical surface in both 5xFAD and WT mice showed increased vessel length, vessel and junction densities. The number of collateral vessels between the middle cerebral artery (MCA) and the anterior and posterior cerebral arteries decreased with age but collateral diameters were significantly increased only in 5xFAD mice. MCA total vessel length and junction density were decreased in 5xFAD mice compared to WT at 4 months. Analysis of ^18^F-FDG cortical uptake revealed significant differences between WT and 5xFAD mice spanning 4-12 months. Broadly, 5xFAD males had significantly increased ^18^F-FDG uptake at 12 months compared to WT mice. In most cortical regions, female 5xFAD mice had reduced ^18^F-FDG uptake compared to WT across their lifespan. While the 5xFAD mouse exhibits AD-like cognitive deficits as early as 4 months of age that are associated with increasing Aβ deposition, we only found significant differences in cortical vascular features in males, not in females. Interestingly, 5xFAD male and female mice exhibited opposite effects in ^18^F-FDG uptake. The MCA supplies blood to large portions of the somatosensory cortex and portions of the motor and visual cortex and increased vessel lengths alongside decreased collaterals coincided with higher metabolic rates in 5xFAD mice. Thus, a potential mismatch between metabolic demand and vascular delivery of nutrients in the face of increasing Aβ deposition could contribute to the progressive cognitive deficits seen in the 5xFAD mouse model.

## 1 Introduction

Alzheimer’s disease (AD) affects 6.7 million American age 65 and older and is the fifth leading cause of death in this population (Alzheimer’s Association, 2023). A hallmark signature in human AD is the progressive cognitive decline and memory deficits that on autopsy are characterized by deposition of extracellular beta-amyloid (Aβ) plaques and intracellular neurofibrillary tangles (Avila and Perry, 2021). Advances in positron emission tomography (PET) using a variety of ligands targeting Aβ and tau proteins have greatly facilitated diagnostic identification of AD (Buckley, 2021). However, less studied is whether vascular alterations precede or contribute to the progression of AD. Recent work suggests that vascular dysfunction contributes to AD pathology (Iturria-Medina et al., 2016) and blood-brain barrier (BBB) disruption (Sweeney et al., 2018). Clinically, cerebral hypoperfusion (Roher et al., 2012; Thomas et al., 2015) has been reported to contribute to hypometabolism (Levin et al., 2021). Morphologically, studies reveal damaged vasculature (Bennett et al., 2018), appearance of string vessels (Hunter et al., 2012), and loss of pericytes (Ding et al., 2020). The bulk of AD-related cerebrovascular studies have focused on cerebral amyloid angiopathy (CAA) that is often associated with cumulative cardiovascular risk factors resulting in diminished brain vascular function (Szidonya and Nickerson, 2023).

Preclinical mouse models of AD have immensely assisted investigations into a mechanistic understanding of the pathophysiology of AD. These studies have provided novel insights and explored potential therapeutics for AD. However, current rodent models of AD (see (Jullienne et al., 2022b)) do not adequately recapitulate the complexity and evolution of human AD. Recent efforts, such as the Model Organism Development & Evaluation for Late-Onset Alzheimer’s Disease (MODEL-AD) consortium (model-ad.org), aim to develop more realistic sporadic mouse models (Oblak et al., 2020). As we have recently reviewed, numerous AD rodent models exhibit altered vascular topology (Szu and Obenaus, 2021). One AD mouse model that has been extensively studied is the 5xFAD mouse (Oblak et al., 2021).

The 5xFAD mouse model has been deeply phenotyped in terms of amyloid burden, Aβ biochemical levels, neuropathology, neurophysiology, and behavior (Forner et al., 2021). It has also been used to probe the effects of amyloidosis (Oblak et al., 2021) on neuronal and glial cells. A small number of studies report abnormal and disconnected capillary segments with reduced junctions and vessel leakage early in life (2-4 months) (Kook et al., 2012; Giannoni et al., 2016; Ahn et al., 2018). These studies noted that in the 5xFAD mouse vascular changes precede frank Aβ deposition and neuronal loss. Alongside these morphological studies, studies have found altered metabolic utilization using ^18^F-fluorodeoxyglucose (FDG) PET. Choi and colleagues reported increased FDG uptake in the hippocampus in aged (8-12 months) 5xFAD mice (Choi et al., 2021). Other studies noted reduced FDG uptake in the brain at 7 months of age of female 5xFAD mice (Bouter et al., 2021) with similar reductions in male mice (Franke et al., 2020). These disparate findings may be related to PET analysis methods, notably standardization approaches.

A gap in the scientific literature is that no studies have phenotyped the vasculature of AD mouse models, specifically the 5xFAD, through their lifespan nor across sex. Most studies use only a single or perhaps two timepoints, as recently reviewed (Szu and Obenaus, 2021). Also, as seen in the neuroscience field, many studies tended to focus predominately on one sex only (Shansky and Woolley, 2016). Our recent neuroimaging review of AD mouse models noted numerous studies in which sex was often not specified (Jullienne et al., 2022b), despite a body of research showing strong sex differences in AD (Udeh-Momoh et al., 2021). Given clinical, and scant preclinical, studies that report cerebrovascular perturbations in AD, we sought to investigate the vascular phenotype of 5xFAD male and female mice and their wild type (WT) littermates across age, from 4 to 12 months, focusing exclusively on the cortical vasculature. We used high resolution magnetic resonance imaging (MRI) to assess brain regional volumes, followed by our vessel painting approach (Salehi et al., 2018) to describe the angioarchitecture in these mice. In a separate cohort of 5xFAD mice, we measured brain glucose metabolism with ^18^F-FDG PET to establish the coupling between perfusion and metabolism. The 5xFAD mouse model of familial AD presents a fast progression of the disease and high expression of Aβ as early as 1.5 months of age, with amyloid deposition and gliosis observed at 2 months. At 9 months, there is a frank amyloid deposition which results in neurodegeneration accompanied by neuronal loss (Oakley et al., 2006). Memory deficits appear at 4 months of age (Oakley et al., 2006; Girard et al., 2014). Here we report on the vascular consequences of aging in male and female 5xFAD mice from 4 through 12 months of age.

## 2 Material and Methods

### 2.1 Animals

All animal experiments and care complied with federal regulations and were approved by the University of California Irvine (UCI) and the Indiana University (IU) Institutional Animal Care and Use Committees. We used the 5xFAD mouse model of Alzheimer’s disease (5xFAD hemizygous, referred to as 5xFAD) and control littermates (C57BL/6J, referred to as WT). All WT and 5xFAD mice used in the present study were provided courtesy of UCI and IU MODEL-AD Consortia. The 5xFAD mice overexpress the APP(695) transgene which harbors the Swedish (K670N, M671L), Florida (I716V), and London (V7171) mutations, and the PSEN1 transgene harboring the M146L and L286V mutations, and were made congenic on the C57BL/6J background to alleviate the concern for allele segregation observed on the hybrid background. Animals were given ad libitum access to food and water, housed with nesting material in a temperature-controlled facility with a 12:12h light/dark cycles. Two cohorts were utilized in this study: 1) UCI: One cohort of animals was used for vessel painting and *ex vivo* MRI experiments (4 groups: males WT, females WT, males 5xFAD and females 5xFAD at 3 timepoints: 4, 8 and 12 months of age, with N=8-9 per group). 2) IU: A second cohort was used for ^18^F-FDG PET experiments (4 groups: males WT, females WT, males 5xFAD and females 5xFAD at 3 similar timepoints: 4, 6 and 12 months of age, with N=9-12 per group, see Table 1). Supplementary Figure 1 outlines the experimental timeline utilized in the present study.

**Table 1:**
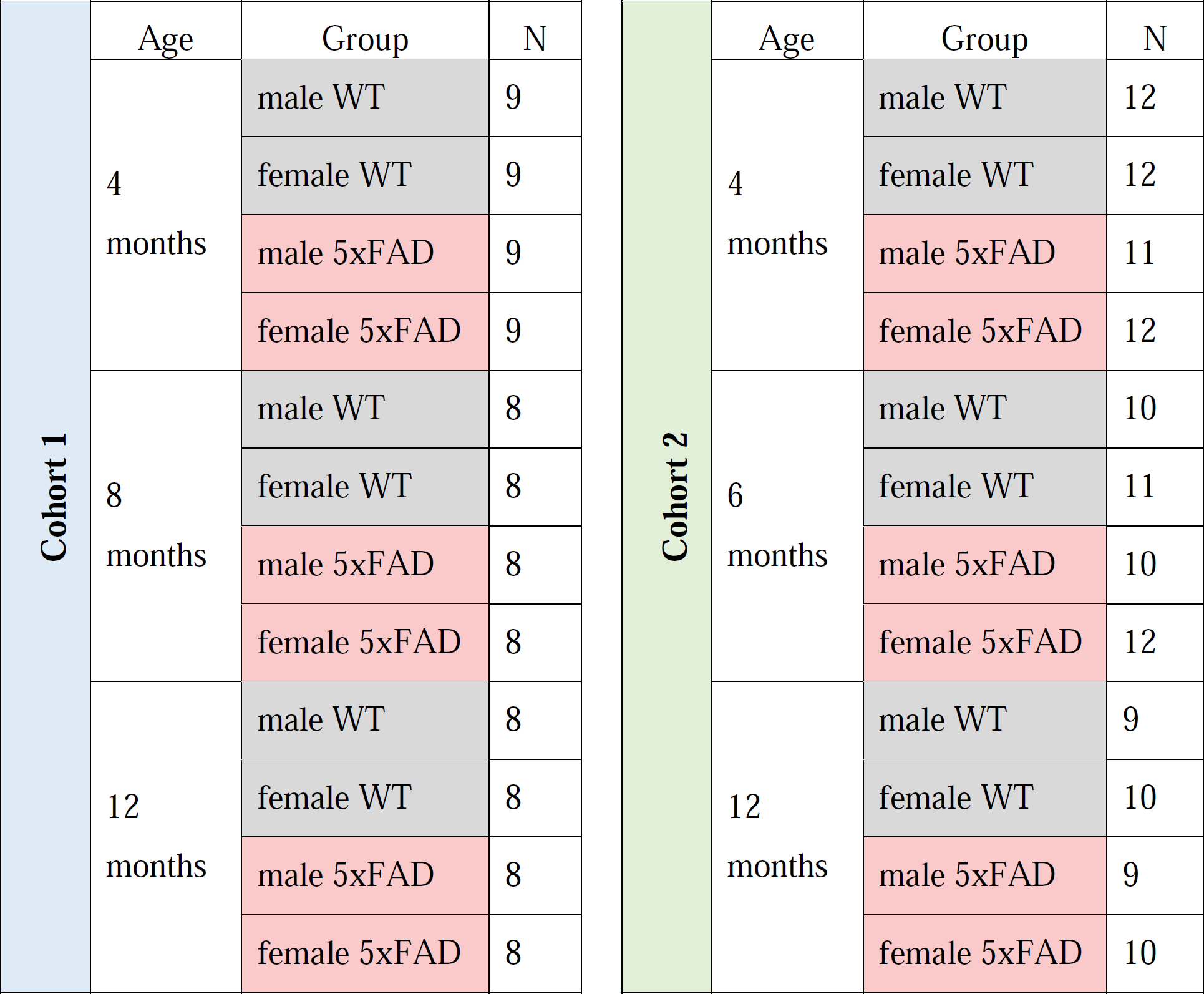
Number of mice per group and in each cohort.

| Cohort 1 | Age | Group | N | Cohort 2 | Age | Group | N |
| --- | --- | --- | --- | --- | --- | --- | --- |
|  | 4 months | male WT | 9 |  | 4 months | male WT | 12 |
|  |  | female WT | 9 |  |  | female WT | 12 |
|  |  | male 5xFAD | 9 |  |  | male 5xFAD | 11 |
|  |  | female 5xFAD | 9 |  |  | female 5xFAD | 12 |
|  | 8 months | male WT | 8 |  | 6 months | male WT | 10 |
|  |  | female WT | 8 |  |  | female WT | 11 |
|  |  | male 5xFAD | 8 |  |  | male 5xFAD | 10 |
|  |  | female 5xFAD | 8 |  |  | female 5xFAD | 12 |
|  | 12 months | male WT | 8 |  | 12 months | male WT | 9 |
|  |  | female WT | 8 |  |  | female WT | 10 |
|  |  | male 5xFAD | 8 |  |  | male 5xFAD | 9 |
|  |  | female 5xFAD | 8 |  |  | female 5xFAD | 10 |

### 2.2 ^18^F-FDG-PET/MRI Methods and Analyses (IU)

Two days prior to positron emission tomography (PET) imaging, mice were anesthetized to prevent movement (induced with 5% isoflurane in 95% medical oxygen, maintained with 1-3% isoflurane). T2-weighted (T2W) MRI images were acquired using a clinical 3T Siemens Prisma MRI scanner (Singo, v7.0), with a 4-channel mouse head coil, bed, and anesthesia system (RapidMR). Post localization, SPACE3D sequences were acquired with the following parameters: TA: 5.5 minutes; TR: 2080ms; TE: 162ms; ETL: 57; FS: On; Average: 2; Excitation Flip Angle: 150; Norm Filter: On; Restore Magnetization: On; Slice Thickness: 0.2 mm; Matrix: 171 × 192; field of view: 35 × 35 mm, yielding 0.18 × 0.18 × 0.2 mm resolution images per our previous work (Kotredes et al., 2021; Oblak et al., 2021; Onos et al., 2022).

PET tracer administration was performed in conscious mice injected intraperitoneally with 3.7 to 9.25 MBq (0.1 to 0.25 mCi) of ^18^F-FDG, and were returned to their warmed home cage for 45 min to permit tracer uptake(Oblak et al., 2021). Post uptake, animals were anesthetized with 5% isoflurane (95% medical oxygen), placed on the imaging bed, landmarked, and scanned on the IndyPET3 scanner (Hutchins et al., 2008). During acquisition, the anesthetic plane was maintained with 1-3% isoflurane (balance medical oxygen). Upon completion, calibrated listmode data were reconstructed into a single-static image with a minimum field of view of 60 mm using filtered-back-projection, and were corrected for decay, random coincidence events, and dead-time loss (Soon et al., 2007) per our previous work (Kotredes et al., 2021; Oblak et al., 2021; Onos et al., 2022).

All PET and MRI images were co-registered using a 9 degrees of freedom ridged-body mutual information-based normalized entropy algorithm (Studholme et al., 1998), and mapped to stereotactic mouse brain coordinates (Franklin and Paxinos, 2013) using Analyze 12 (AnalyzeDirect, Stilwell KS, RRID:SCR_005988). Post-registration, 56 bilateral brain regions were extracted, left and right regions averaged, and ratioed to the cerebellum yielding 27 unique specific uptake value ratios (SUVR) volumes of interest. Regional data were summarized in Excel. Cortical regions mimicking the axial surface of vessel painted brains were utilized for comparative analyses (see Figure 9A).

### 2.3 Vessel Painting Methods and Analyses

The vessel painting technique is based on the ability of the fluorescent dye 1,1’-dioctadecyl-3,3,3’3’-tetramethylindocarbocyanine perchlorate (DiI, Life Technologies, Carlsbad, CA, USA) to bind to lipid membranes. The protocol was modified from previous studies (Hughes et al., 2014; Salehi et al., 2018). Mice were intraperitoneally injected with heparin and sodium nitroprusside and 5 minutes later, were anesthetized with an intraperitoneal injection of ketamine (90 mg/kg) and xylazine (10 mg/kg). Vessel painting was performed by manually injecting a solution of DiI (0.3 mg/mL in PBS containing 4% dextrose, total volume of 500 µL) into the left ventricle, followed by a 10 mL PBS flush and a 20 mL 4% PFA perfusion using a peristaltic pump (8.4 mL/min). Heads were post-fixed in 4% PFA for 24 hours, rinsed in PBS and stored at 4°C in PBS until magnetic resonance imaging (MRI). Successfully vessel painted brains were selected if they showed uniform pink staining and excellent staining of large and small vessels on the cortical surface. In this study, 84% (84/100) of the brains were successfully stained and analyzed.

After *ex vivo* MRI, brains were extracted from the skull and were imaged using a wide-field fluorescence microscope (Keyence BZ-X810, Keyence Corp, Osaka, Japan). Axial images of the entire brain were acquired at 2X magnification using the Z-stack feature (∼42 images, step size 25.2 µm). A portion of the left middle cerebral artery (MCA) was also imaged for each sample using a confocal microscope (10X magnification, Z-stack 30 images, step size 1.51 μm, Olympus FV3000, Olympus Scientific).

All vascular analysis methods utilized in the current study have been previously published (Salehi et al., 2018; Jullienne et al., 2022a). Briefly, classical vessel analysis was performed using AngioTool Software (RRID:SCR_016393) for measures of vessel density, length, and number of junctions (Zudaire et al., 2011). The ImageJ plugin “FracLac” (ImageJ, RRID:SCR_003070) was used to analyze vascular complexity using fractal analysis (Karperien et al., 2013). Collateral vessels provide an alternate route of blood perfusion in case of occlusion and have been shown to decrease with aging (Faber et al., 2011). The number of collateral vessels between MCA and posterior and anterior cerebral arteries (PCA and ACA) were manually counted for each brain. Collateral diameters were measured using ImageJ. Areas of blood-brain barrier leakage evidenced by DiI extravasation were analyzed from 10X images using ImageJ. All data were extracted and summarized in MS Excel.

### 2.4 Ex vivo Magnetic Resonance Imaging (MRI) (UCI)

Brain imaging was conducted on *ex vivo* in-skull brains on either a Bruker Biospec UltraShield Refrigerated (USR) 9.4T magnet or a Bruker Avance Instrument 11.7T magnet. Samples underwent T2-weighted 3D rapid-acquisition with relaxation enhancement (T2 RARE) imaging for regional and total brain volumes. Table 2 lists the imaging parameters. Brain tissue was extracted from the T2 RARE scans using 3D Pulse-Coupled Neural Networks (PCNN3D) in MATLAB R2017a (RRID:SCR_001622). Extraction masks were then reviewed and adjusted using ITK-SNAP (version 3.8.0, RRID:SCR_002010, (Yushkevich et al., 2006)). After extraction, scans underwent N4 Bias field correction (Tustison et al., 2010). Next, a modified version of the Australian Mouse Brain Mapping Consortium Atlas (AMBMC, (Richards et al., 2011)) was fit to each animal and regional labels were applied with Advanced Normalization Tools (ANTS, RRID:SCR_004757). Whole brain, cerebrum, and regional (40 bilateral regions) volumes were extracted and exported to Microsoft Excel for summary analysis.

**Table 2:**
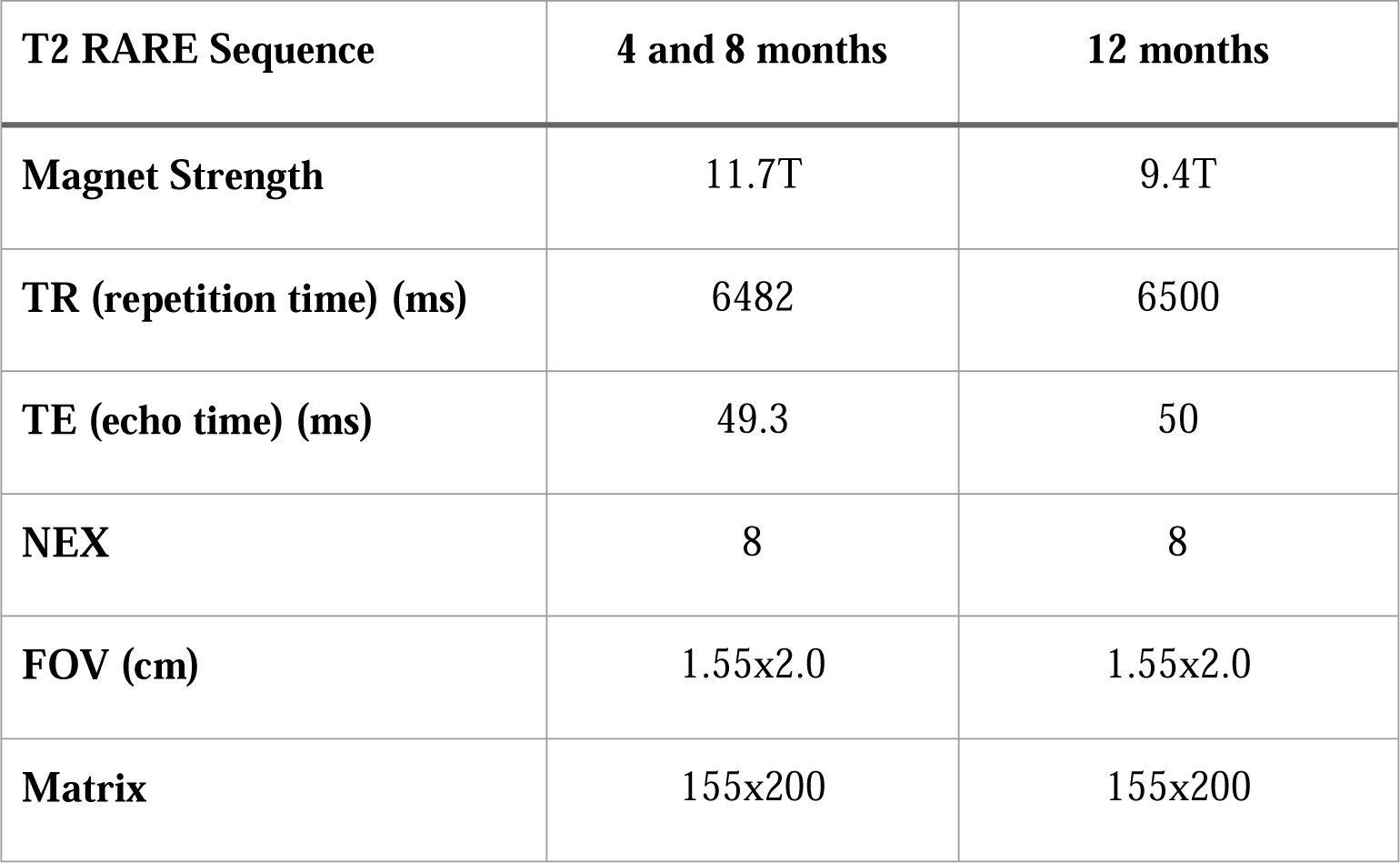
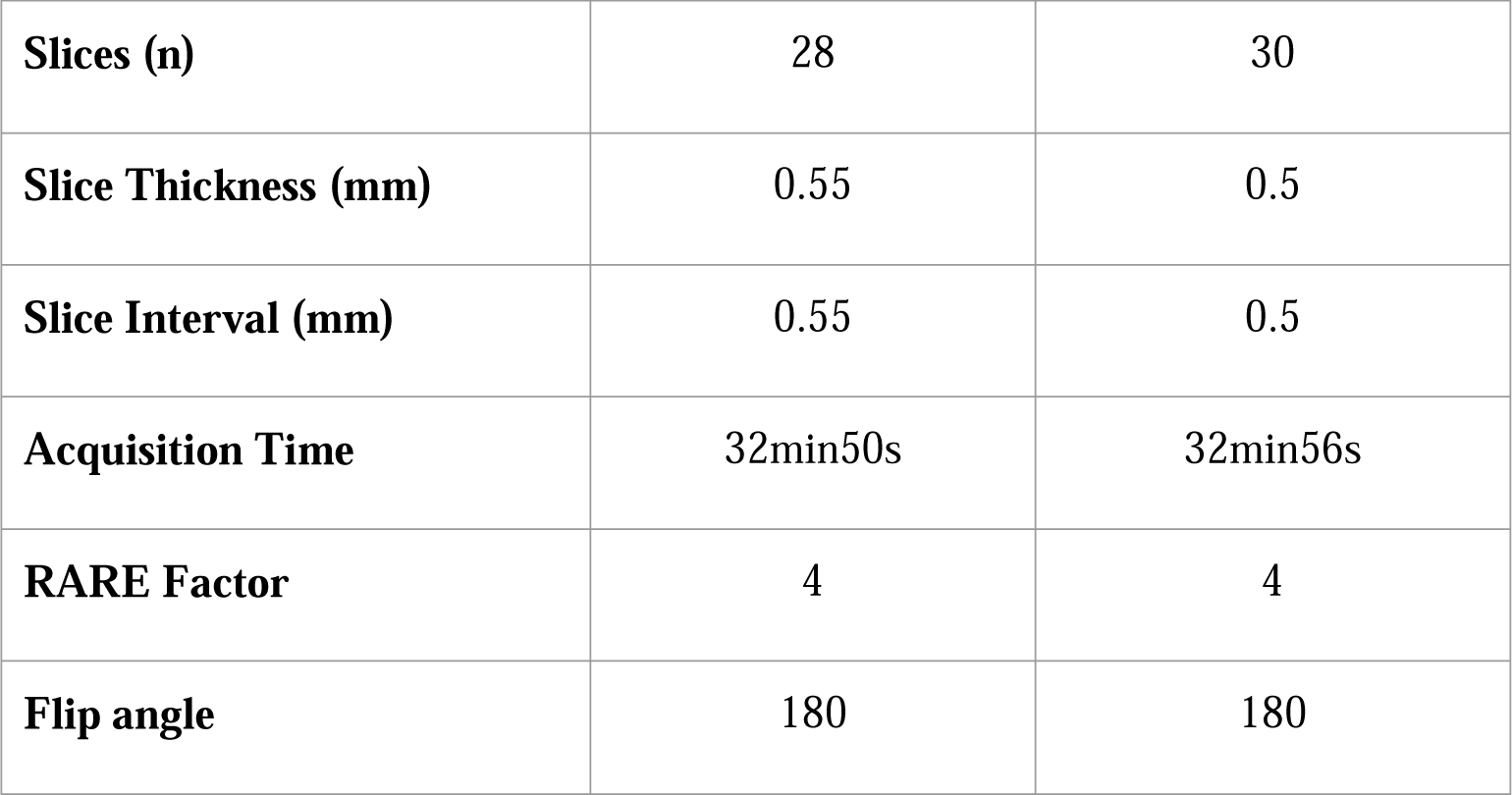
Magnetic Resonance Imaging parameters for Cohort 1.

### 2.5 Statistical Analyses

All data from each cohort underwent curation and validation. Data were assessed for outliers using interquartile range prior to statistical testing. GraphPad Prism 9 software (GraphPad, RRID:SCR_002798) was used to perform statistical analysis. Two-way ANOVA and Sidak’s multiple comparisons post-hoc testing was utilized for vessel analysis, cerebrum, whole brain, regional brain volumes and brain metabolism. For the DiI extravasation analysis, we used non-parametric tests (Kruskal-Wallis) as the number of replicates were smaller in some groups. Graphs are presented with post-hoc statistical significance noted as \**p*<0.05, \*\**p*<0.01, \*\*\**p*<0.001, or **** *p*<0.0001. For two-way ANOVA, effect of age is shown as δ *p*<0.05, δδ *p*<0.01, δδδ *p*<0.001, or δδδδ *p*<0.0001, and effect of genotype as # *p*<0.05, ## *p*<0.01, ### *p*<0.001, or #### *p*<0.0001. In the results, graphs appear as box and whisker plots (denoting median, lower and upper quartile, maximum and minimum) or line graphs presented as mean ± SEM. Age and genotype statistical findings are reported above each data type in the figures.

## 3 Results

### 3.1 Cerebrum brain volumes

High-resolution MRI scans of 5xFAD and WT mice were undertaken at each age (4, 8, and 12 months, Figure 1A). Cerebrum volumes significantly increased with age in both WT and 5xFAD mice (two-way ANOVA, δδδδ p<0.0001, Figure 1B), independent of the genotype. A genotype effect was detected (two-way ANOVA, # p=0.024, Figure 1B) which was driven by females. There was a significant decrease in cerebrum volume at 8 months of age between WT and 5xFAD females from 351.60±4.97 to 333.81±4.10mm^3^ (* p=0.038, Figure 1C). The cerebrum volumes of 8-month-old WT males were significantly lower than WT females at the same age (** p=0.009, Figure 1C).

**Figure 1:**
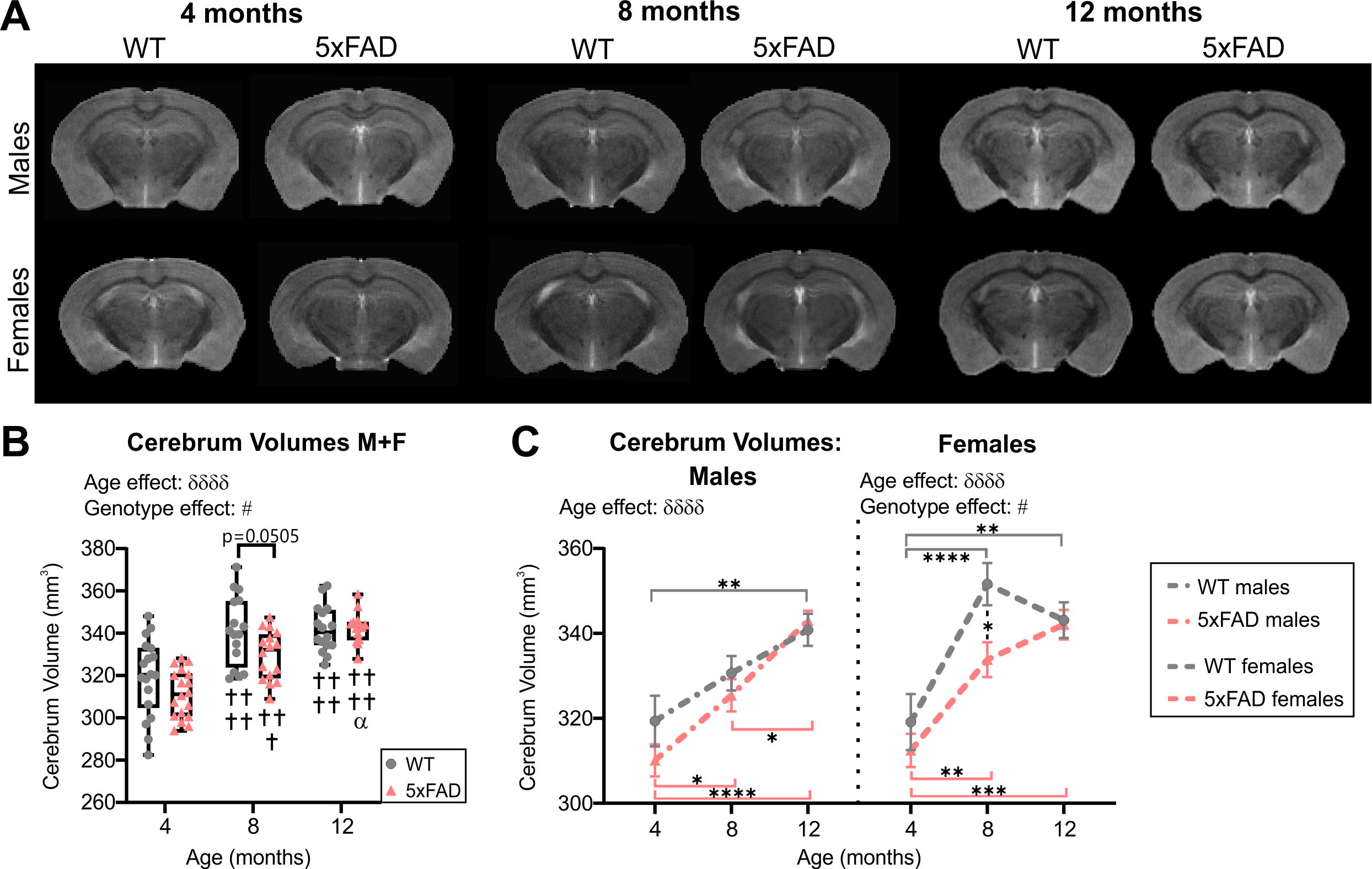
Cerebrum volumes across age. **(A)** Representative T2 RARE images at Bregma level -2.1mm for male and female WT and 5xFAD mice across age. **(B)** Cerebrum volumes increased with age in WT and 5xFAD mice but more rapidly in WT mice. **(C)** Cerebrum volumes for males and females revealed a significant difference between genotypes at 8 months only in females. δ reports a significant effect of age (two-way ANOVA, δδδδ=p<0.0001), # shows a significant effect of the group/genotype (two-way ANOVA, #=p<0.05); for multiple comparisons, † shows a significant difference from the 4 months timepoint and α a significant difference from 8 months timepoint (α=p<0.05, †††=p<0.001, ††††=p<0.0001), *=p<0.05, **=p<0.01, ***=p<0.001, ****=p<0.0001.

### 3.2 Cortical vascular topology: Genotype differences

Wide-field fluorescent microscopy of the axial cortical surface allows visualization of vessels up to a depth of 1 mm. Classical vessel features (density, length, junctions) of the cortical vascular network revealed no overt genotype effects for any of the metrics studied (two-way ANOVA, vessel density p=0.825, junction density p=0.739, total vessel length p=0.788, average vessel length p=0.092). A significant effect of age was observed in both WT and 5xFAD for vessel density (two-way ANOVA, δδ p=0.001, Figure 2A, 2E), junction density (two-way ANOVA, δδδδ p<0.0001, Figure 2B, 2F), total vessel length (two-way ANOVA, δδδδ p<0.0001, Figure 2C, 2G), and average vessel length (two-way ANOVA, δδδδ p<0.0001, Figure 2D,2H). These four vessel metrics were increased with age in both groups. Junction density was significantly increased between 4 and 12 months for WT (Sidak’s test, *** p=0.0008) and 5xFAD mice (Sidak’s test, * p=0.025, Figure 2B, 2F), total vessel length was significantly increased between 4 and 12 months for WT (Sidak’s test, ** p=0.008) and 5xFAD mice (Sidak’s test, ** p=0.007, figure 2C and 2G), and average vessel length was significantly increased between 4 and 12 months for WT (Sidak’s test, **** p<0.0001) and 5xFAD mice (Sidak’s test, * p=0.037, Figure 2D, 2H). These results suggest a continuous growth/maturation of vessels during aging in both WT and transgenic mice independent of genotype.

**Figure 2:**
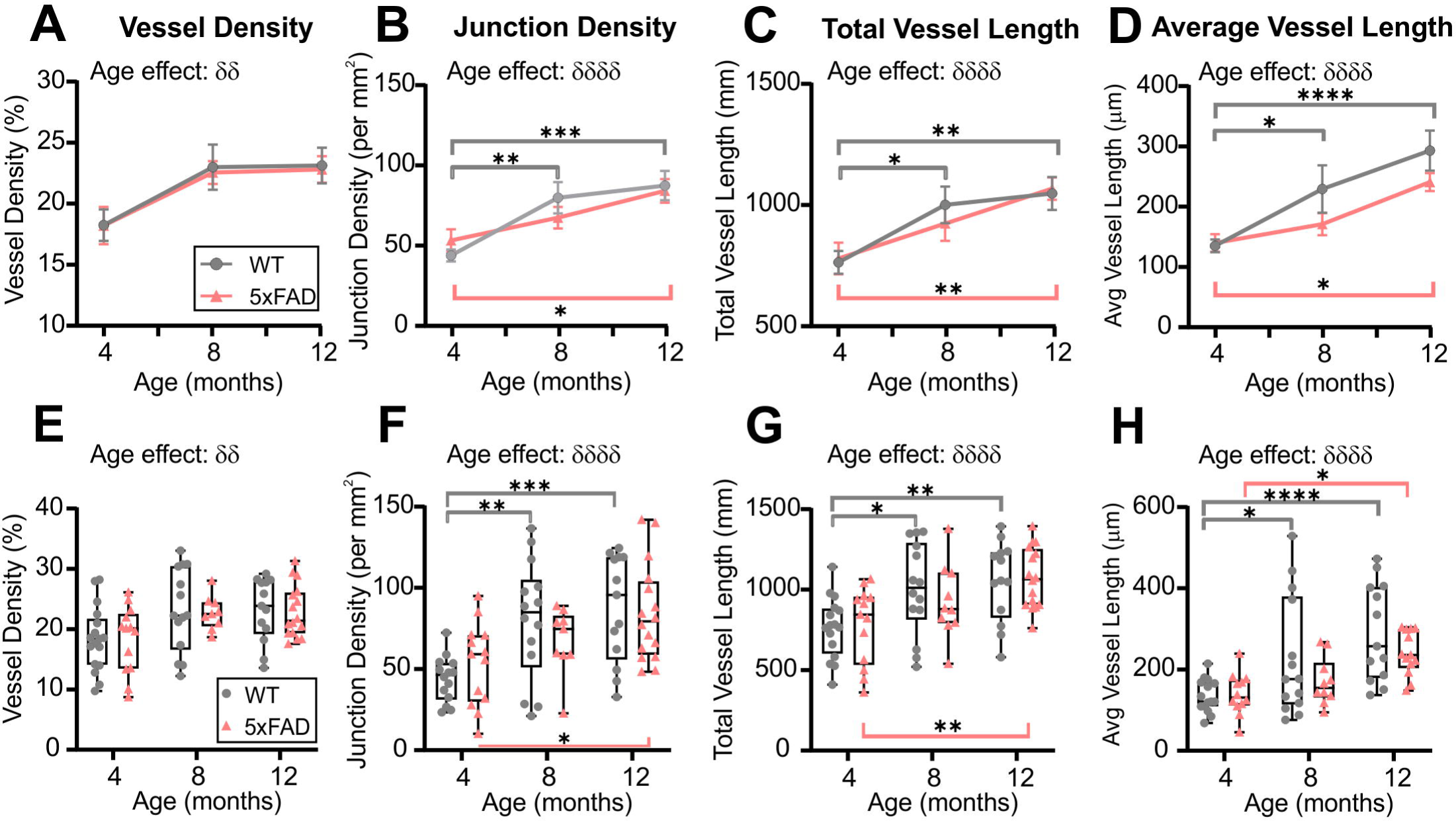
Angioarchitecture of the cortical surface in WT and 5xFAD mice with increasing age. Vessel density **(A, E)**, junction density **(B, F)**, total vessel length **(C, G)**, and average vessel length **(D, H)** are shown for WT and 5xFAD mice at 4, 8, and 12 months. Within each genotype, there were significant increases in vascular measures with age. No significance between genotypes was observed. δ shows a significant effect of age (two-way ANOVA, δδ=p<0.01, δδδδ=p<0.0001); for multiple comparisons (Sidak’s test): *=p<0.05, **=p<0.01, ***=p<0.001, ****=p<0.0001, grey line compares WT while red line compares 5xFAD mice.

### 3.3 Cortical vascular topology: Sex differences

When male and female mice were dichotomized, we observed an effect of age clearly driven by males (Figure 3). No effect of age was detected in female mice (two-way ANOVA: p=0.620 for vessel density, p=0.627 for junction density, p=0.315 for total vessel length, p=0.290 for average vessel length). In males, the effect of age was highly significant for all the cortical vessel metrics (two-way ANOVA: δδδ p=0.0005 for vessel density, δδδ p=0.0001 for junction density, δδδ p=0.0001 for total vessel length, δδδδ p<0.0001 for average vessel length). In 5xFAD male mice a significant increase was observed between 4 and 12 months for vessel density (Sidak’s test, ** p=0.004, Figure 3A), junction density (Sidak’s test, ** p=0.001, Figure 3B), total vessel length (Sidak’s test, ** p=0.001, Figure 3C), and average vessel length (Sidak’s test, **** p<0.0001, Figure 3D) (Figure 3I, 3J). Between 4 and 12 months, WT males exhibited a significant increase in vessel density (Sidak’s test, * p=0.050, Figure 3A), junction density (Sidak’s test, * p=0.028, Figure 3B), total vessel length (Sidak’s test, * p=0.025, Figure 3C) and average vessel length (Sidak’s test, ** p=0.002, Figure 3D). No significant differences were observed in female mice in any of the four metrics (Figure 3E-H). These results demonstrate that vascular networks evolve differently with increasing age between males and females, but genotype does not seem to alter features of the cortical vessel network.

**Figure 3:**
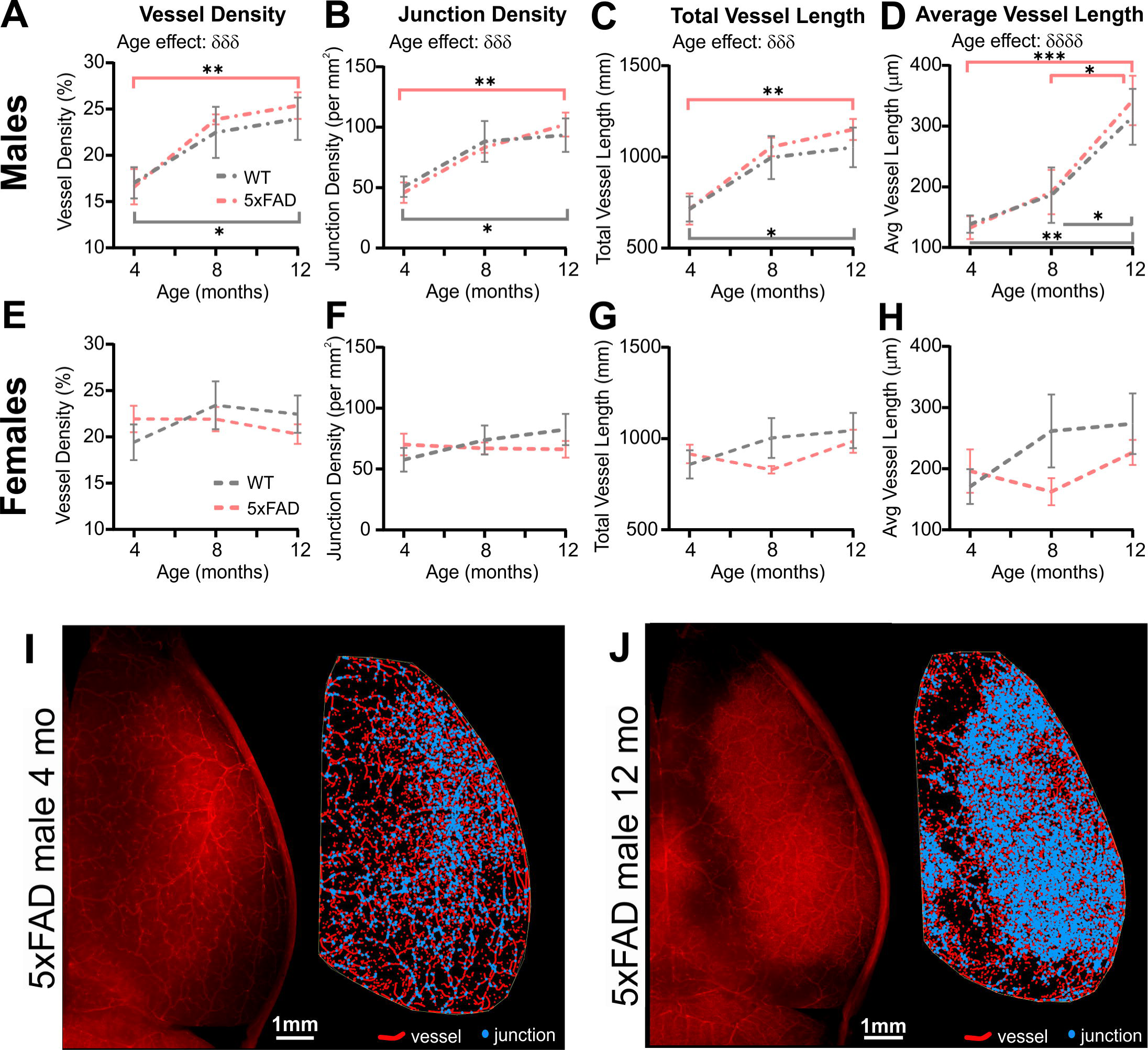
Vascular network features of the cortical surface of male and female WT and 5xFAD mice. Vessel density **(A, E)**, junction density **(B, F)**, total vessel length **(C, G)**, and average vessel length **(D, H)** are shown for male and female WT and 5xFAD mice at 4, 8, and 12 months. The age effect reported in Figure 2 is driven primarily by males as female vascular features were relatively stable. **(I, J)** Representative images of vessel painted hemispheres (left) with Angiotool resultant image (right) in a 4-month-old 5xFAD male **(I)** and a 12-month-old 5xFAD male **(J)** illustrate the increasing junctions. δ reports a significant effect of age (two-way ANOVA, δδδ=p<0.001, δδδδ=p<0.0001); for multiple comparisons (Sidak’s test): *=p<0.05, **=p<0.01, ***=p<0.001.

### 3.4 Cortical vascular complexity

Fractal analysis for complexity of the cortical vessel network in WT and 5xFAD mice across ages was assessed. Fractal histograms with increased maximal frequency reflect increased numbers of vessels whilst a rightward shift of the local fractal dimension (LFD) distribution reports increased vascular complexity (increased LFD). Fractal histogram shape features such as skewness and kurtosis were not altered by age nor genotype (Supplementary Figure 2). Consistent with results obtained from our classical vascular topology analyses (Figures 2, 3), a trend in increased maximum frequency value (ie. increased number of vessels) was observed with age in WT and 5xFAD mice (two-way ANOVA, p=0.064, Supplementary Figure 2D, Figure 4A-C). A significant effect of genotype was also found for the maximal LFD value (two-way ANOVA, ### p=0.0003) with differences in cortical vessel complexity observed at 8 months between WT and 5xFAD mice (Sidak’s test, *** p=0.0004, Supplemental Figure 2C, Figure 4B). Interestingly, this complexity difference was driven by females (Figure 4H), as reflected by the significant increase in maximal LFD value in 5xFAD mice compared to WT at 8 months (two-way ANOVA, ## p=0.005, Sidak’s test **** p<0.0001, Supplementary Figure 2K). In summary, vascular complexity increases with age but more so in female 5xFAD mice.

**Figure 4:**
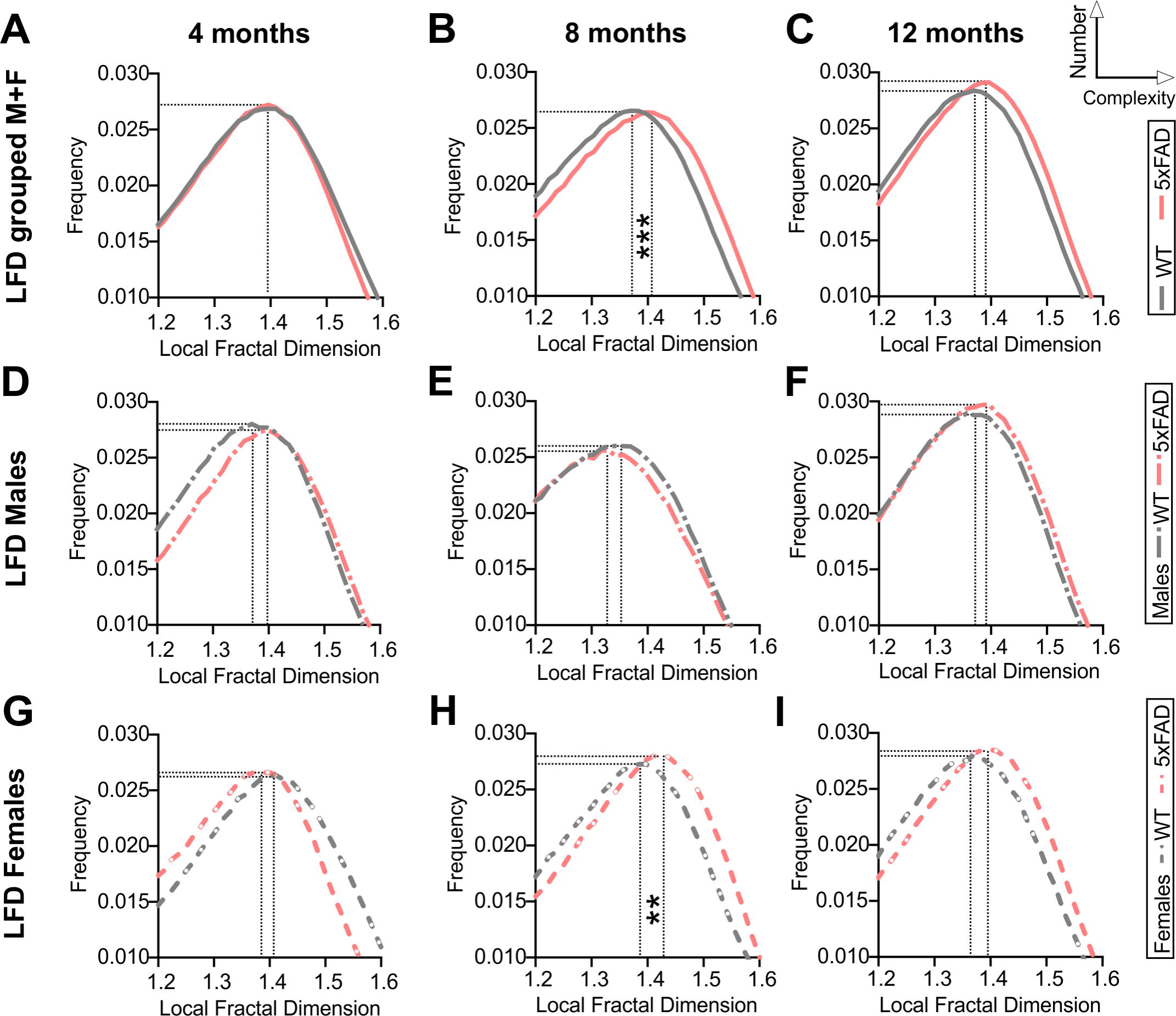
Fractal analysis of vascular network complexity of the cortical surface in WT and 5xFAD mice across ages. Local Fractal Dimension (LFD) histograms for WT and 5xFAD mice at 4 **(A)**, 8 **(B)** and 12 months of age **(C)**. There is a predominant rightward shift in the LFD at 8 and 12 months of age indicating more vascular complexity in 5xFAD mice. LFD histograms are shown for males **(D, E, F)** and females **(G, H, I)**. The maximum frequency value was increased with age in both groups while an effect of genotype was only apparent at 8 months, mainly in females **(H)**. For multiple comparisons across ages and between genotypes (Sidak’s test): **=p<0.01, ***=p<0.001, ****=p<0.0001, based on max LFD value, see Supplementary Figure 2.

### 3.5 Vascular topology of the middle cerebral artery

The global hemispheric cortical vascular network increases with advancing age in both 5xFAD and WT mice, but we then undertook a deeper examination of the middle cerebral artery (MCA) vascular features. Using confocal microscopy, we confirmed a significant age effect in junction density (two-way ANOVA, δδ p=0.004) and total vessel length (δ p=0.028). In general, MCA vessel characteristics in 5xFAD were globally decreased compared to WT mice. Vessel density exhibited a trending decrease (two-way ANOVA, p=0.072, Figure 5A, E). There was a significant genotype effect for junction density (two-way ANOVA, # p=0.021, Figure 5B, F), total vessel length (two-way ANOVA, ## p=0.004, Figure 5C, G) and average vessel length (two-way ANOVA, ### p=0.0002, Figure 5D, H).

**Figure 5:**
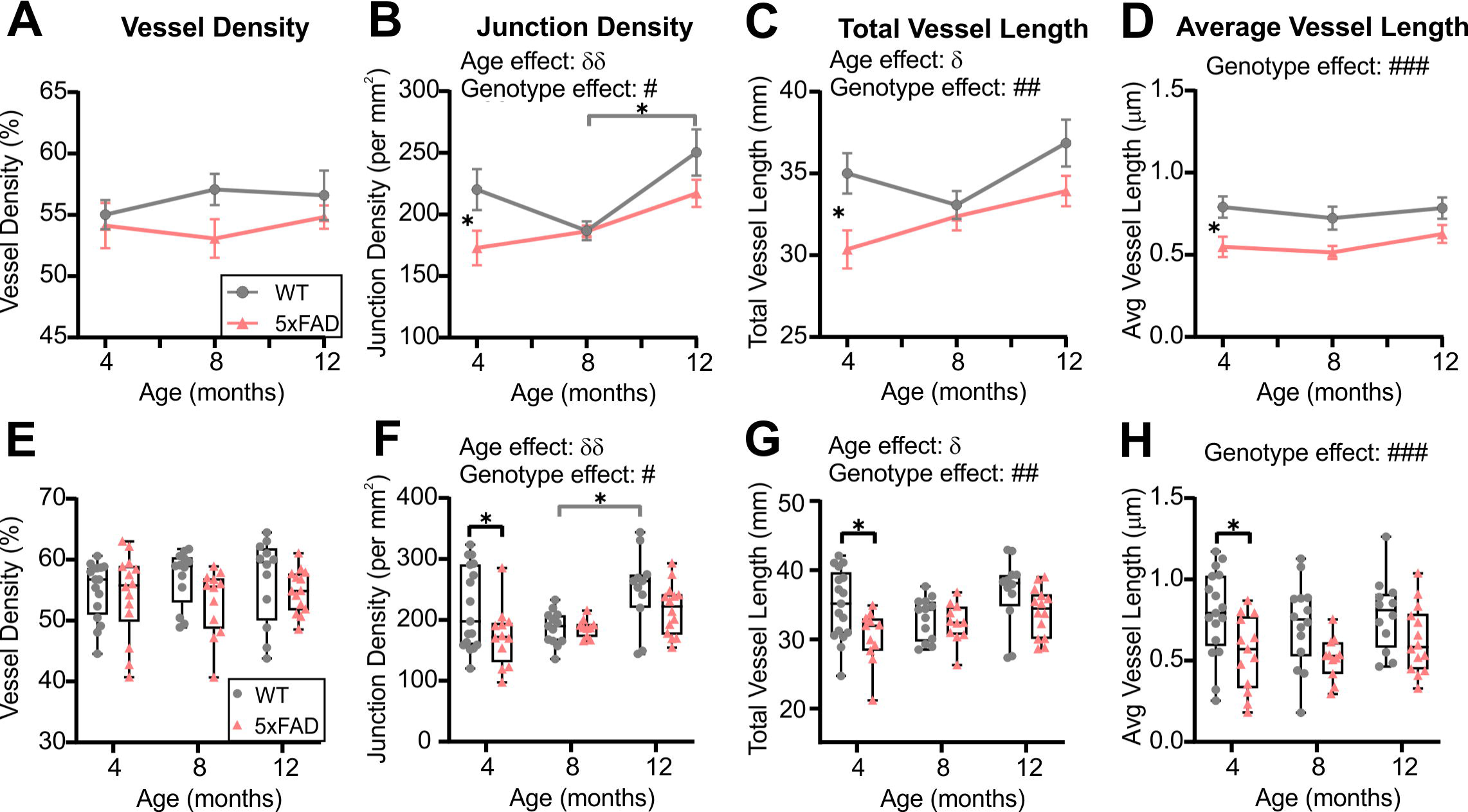
Vessel characteristics of the middle cerebral artery (MCA) in WT and 5xFAD mice across ages (males and females grouped). Vessel density at the level of the M3 MCA branch **(A, E)** showed no temporal age-related changes. Junction density **(B, F)**, total vessel length **(C, G)**, and average vessel length **(D, H)** were assessed at 4, 8, and 12 months of age and exhibited a genotype effect where decreases were observed in 5xFAD compared to the WT mice. δ shows significant effect of age (two-way ANOVA, δ=p<0.05, δδ=p<0.01); # shows a significant effect of genotype (two-way ANOVA, #=p<0.05, ##=p<0.01, ###=p<0.001); for multiple comparisons across ages and between genotypes (Sidak’s test): *=p<0.05.

When mice were dichotomized by sex, we noted that the age and genotype effects were once again driven by males (Figure 6, see also Figure 3). No significant age or genotype effects were detected in females despite an overall decrement in vascular MCA features (Figure 6E-H). In 5xFAD males, there was a significant reduction in junction density (two-way ANOVA, # p=0.062, Figure 6B), total vessel length (## p=0.002, Figure 6C), and average vessel length (# p=0.017, Figure 6D). Thus, the vascular features of the MCA in 5xFAD mice, in particular males, were reduced compared to WT mice. A physiological consequence of these genotypic reductions would be altered nutrient supply and waste removal in vulnerable 5xFAD mice, particularly males.

**Figure 6:**
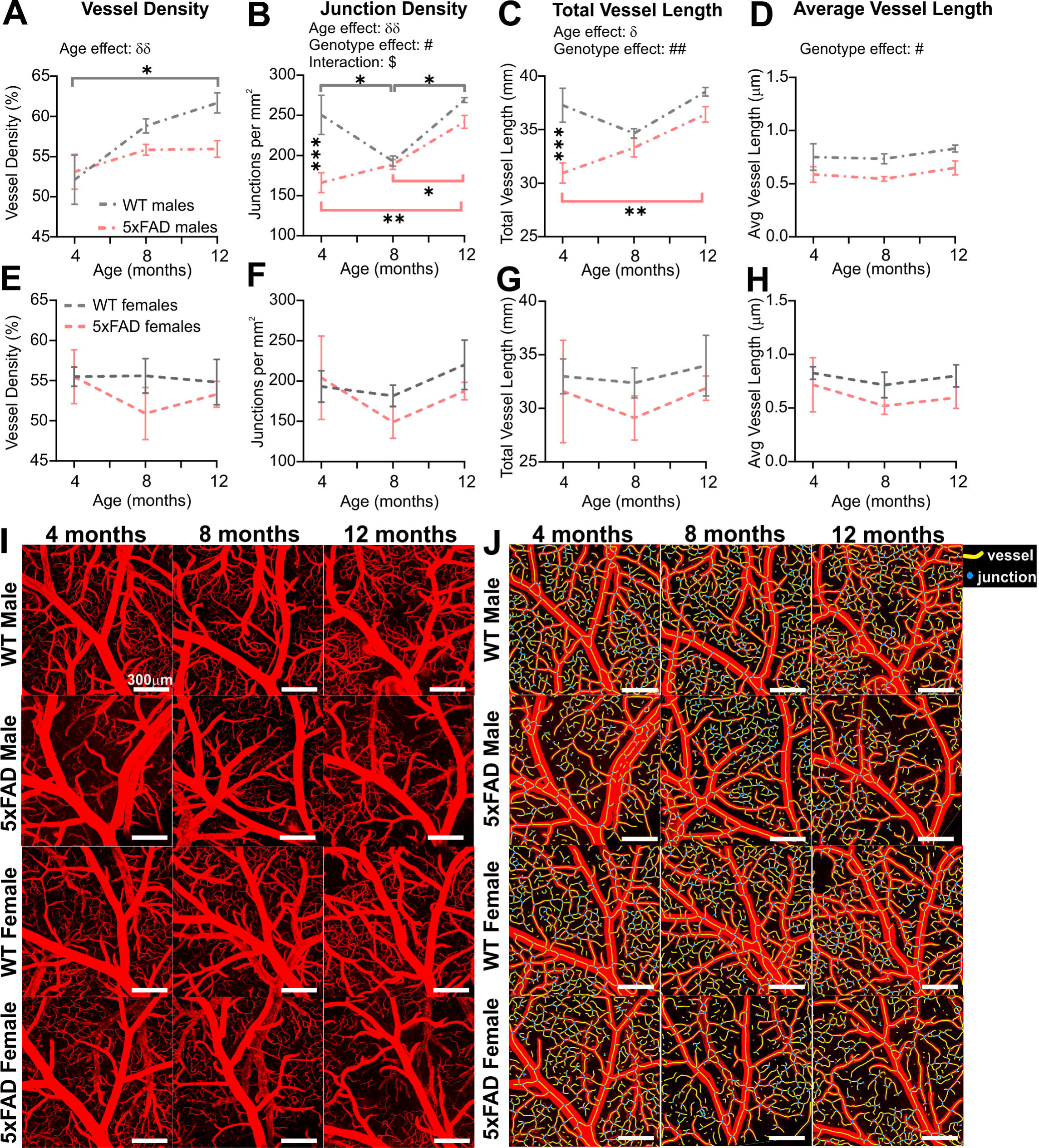
Sex differences of the M3 branch of the middle cerebral artery (MCA) in WT and 5xFAD mice. Vessel density, junction density, total vessel length, and average vessel length were assessed at 4, 8, and 12 months of age in males **(A-D)** and in females **(E-H)**. Significant differences appear to be driven by males. Representative confocal pictures of the MCA portion are shown for male and female mice **(I)**, with the corresponding Angiotool results images **(J)**. δ shows a significant effect of age (two-way ANOVA, δ=p<0.05, δδ=p<0.01); # shows a significant effect of the genotype (two-way ANOVA, #=p<0.05, ##=p<0.01); $ shows a significant interaction between age and genotype (two-way ANOVA, $=p<0.05); for multiple comparisons across time and between genotypes (Sidak’s test): *=p<0.05, **=p<0.01, ***=p<0.001. Scale bar in **I-J** = 300μm.

### 3.6 Blood-brain barrier leakage

Regions of DiI extravasation were first observed in the older animals on confocal images of the MCA (Figure 7A) which prompted a more quantitative analysis. In WT group, 5.9% of mice exhibited DiI extravasation at 4 months whereas 13.3% of 5xFAD mice had leakages. At 8 months, no WT mice had DiI extravasations, however 50% of the 5xFAD mice had leakages. By 12 months, 54.5% of the WT mice and 66.7% of the 5xFAD mice presented with areas of DiI extravasation (Figure 7B). Interestingly the proportion of male and female 5xFAD mice exhibiting leakages were similar at 12 months of age (66.7%) but male WT mice exhibited increased numbers of mice with leakages (75.0%) compared to female WT mice (42.9%). The area encompassing each leak was quantified and at 12 months of age 5xFAD mice demonstrated a significant decrease compared to 8-month-old 5xFAD mice (Kruskal-Wallis test, ** p=0.001). This difference was driven by females (Kruskal-Wallis test, ** p=0.009 for females and p=0.251 for males, Figure 7C). Leakage metrics were not correlated to any of the classical vessel metrics (Supplementary Figure 3).

**Figure 7:**
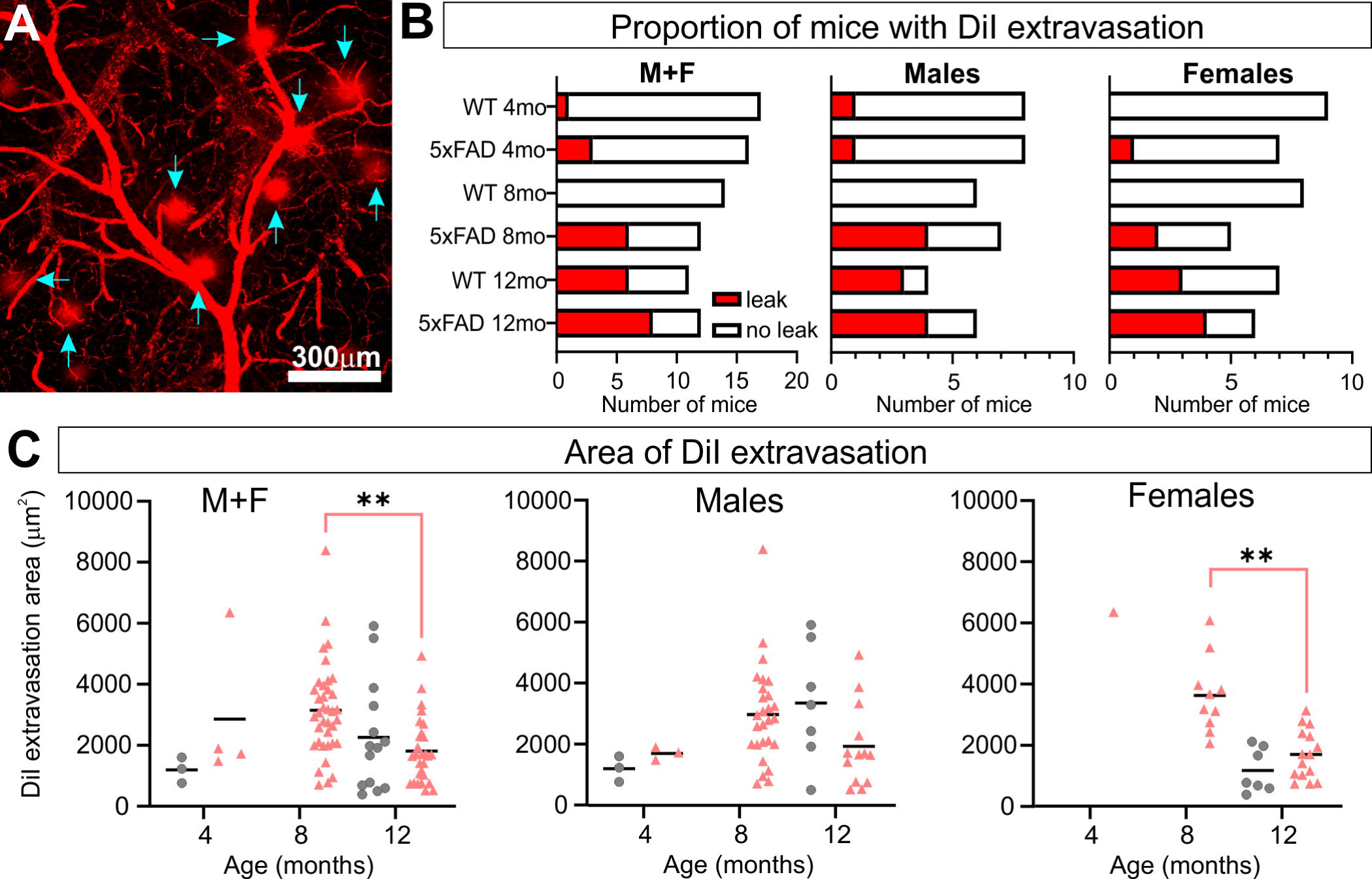
5xFAD mice exhibit increased vascular leakage consistent with blood-brain barrier (BBB) disruption. Representative confocal image of DiI extravasation areas (blue arrows) in an 8-month-old 5xFAD male **(A)**. The proportion of mice with BBB disruptions increased with time in 5xFAD mice relative to WT **(B)** with females having a consistent increasing leak trajectory. Leak area was compiled **(C)** and revealed a maximal increase in leak area at 8 months followed by a reduction at 12 months in 5xFAD mice. For multiple comparisons between genotypes and across time (non-parametric Kruskal-Wallis test): **=p<0.01.

### 3.7 Watershed collaterals

MCA, anterior cerebral artery (ACA) and posterior cerebral artery (PCA) are linked together by a collateral vascular network on the cortical surface, a region of known vulnerability denoted as the watershed territory (Figure 8A). We found a significant decrease in the number of collaterals with advancing age in both WT and 5xFAD mice (two-way ANOVA, δδδδ p<0.0001, Figure 8B, C). Specifically, collaterals were significantly decreased in both WT and 5xFAD mice between 4 and 12 months, and between 8 and 12 months (Figure 8B). No sex differences were observed in collateral numbers. We also assessed collateral vessel diameters, which exhibited an age and a genotype effect (two-way ANOVA, age: δδ p=0.001, genotype: ## p=0.005, Figure 8D, E). The vessel diameter age effect was significant in males (two-way ANOVA, δδ p=0.001, Figure 8F) and trending in females (two-way ANOVA, p=0.061, Figure 8G). The significant genotype effect was in females (two-way ANOVA, # p=0.044), with a difference between WT and 5xFAD females at 8 months (Sidak’s test, * p=0.027, Figure 8G), but only trending in males (two-way ANOVA, p=0.068, Figure 8F).

**Figure 8:**
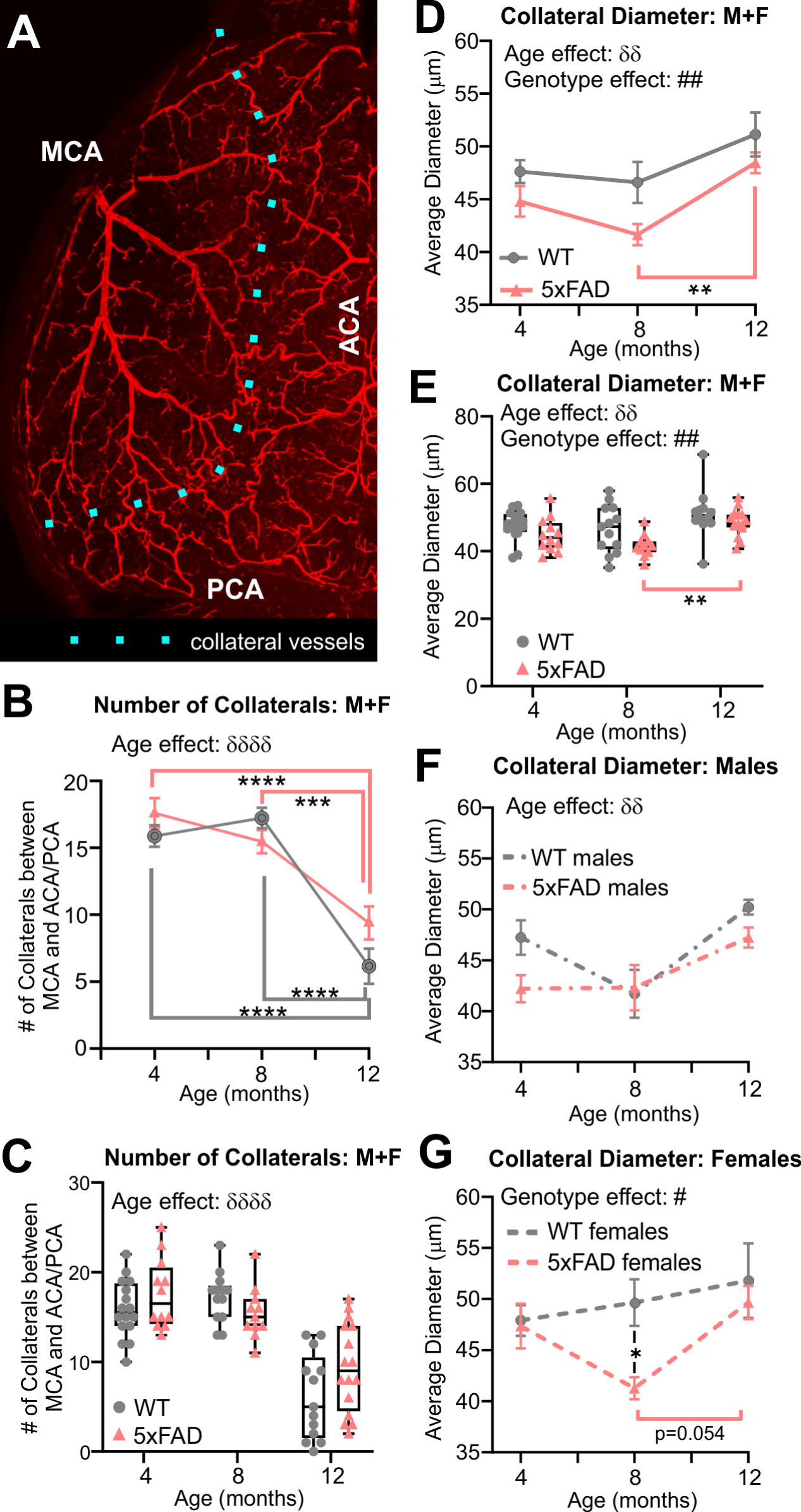
Quantitative vascular assessment of the watershed region. The number of collaterals vessels at the intersection of the MCA, ACA and PCA were counted along the dotted line **(A)**. Collateral vessels decreased with age in both groups independent of genotype, particularly at 12 months of age in both WT and 5xFAD mice **(B, C)**. Average diameter of collateral vessels was reduced in 5xFAD mice compared to WT **(D, E)**. Average collateral diameters exhibited different temporal progression in males **(F)** compared to females **(G)**, especially at 8 months. δ shows an effect of genotype (two-way ANOVA, δδ=p<0.01, δδδδ=p<0.0001), # shows an effect of the genotype (two-way ANOVA, ##=p<0.01), for multiple comparisons across ages and between genotypes (Sidak’s test) **=p<0.01, ***=p<0.001, ****=p<0.0001.

### 3.8 Metabolic perturbations

Metabolic changes within 5xFAD and WT mice across age (4, 6, and 12 months) were assessed using ^18^F-FDG-PET (Figure 9A, B). Cerebral cortex ^18^F-FDG measurements that complemented our axial cortical surface vessel topology analyses were compared. No cortical metabolic differences between the genotypes were observed at 4 months of age (Figure 9C); similarly, no overt metabolic changes at 4, 6 or 12 months of age were found when male and female mice were combined (except decreased ^18^F-FDG uptake in 5xFAD mice in visual cortex at 6 months). When the genotypes were dichotomized by sex, 5xFAD males at 12 months had significant increased uptake of ^18^F-FDG in the secondary motor cortex (M2), the retrosplenial dysgranular cortex (RSC) and the primary/secondary visual (V1/V2) cortex compared to WT (t-tests, M2: * p=0.017, RSC: ** p=0.002, V1/V2: ** p=0.009, Figure 9C, Supplementary Figure 4). In 5xFAD females there was greater variability, with glucose metabolism significantly decreased in the visual area compared to WT (t-test, * p=0.036, Figure 9C, Supplementary Figure 4).

**Figure 9:**
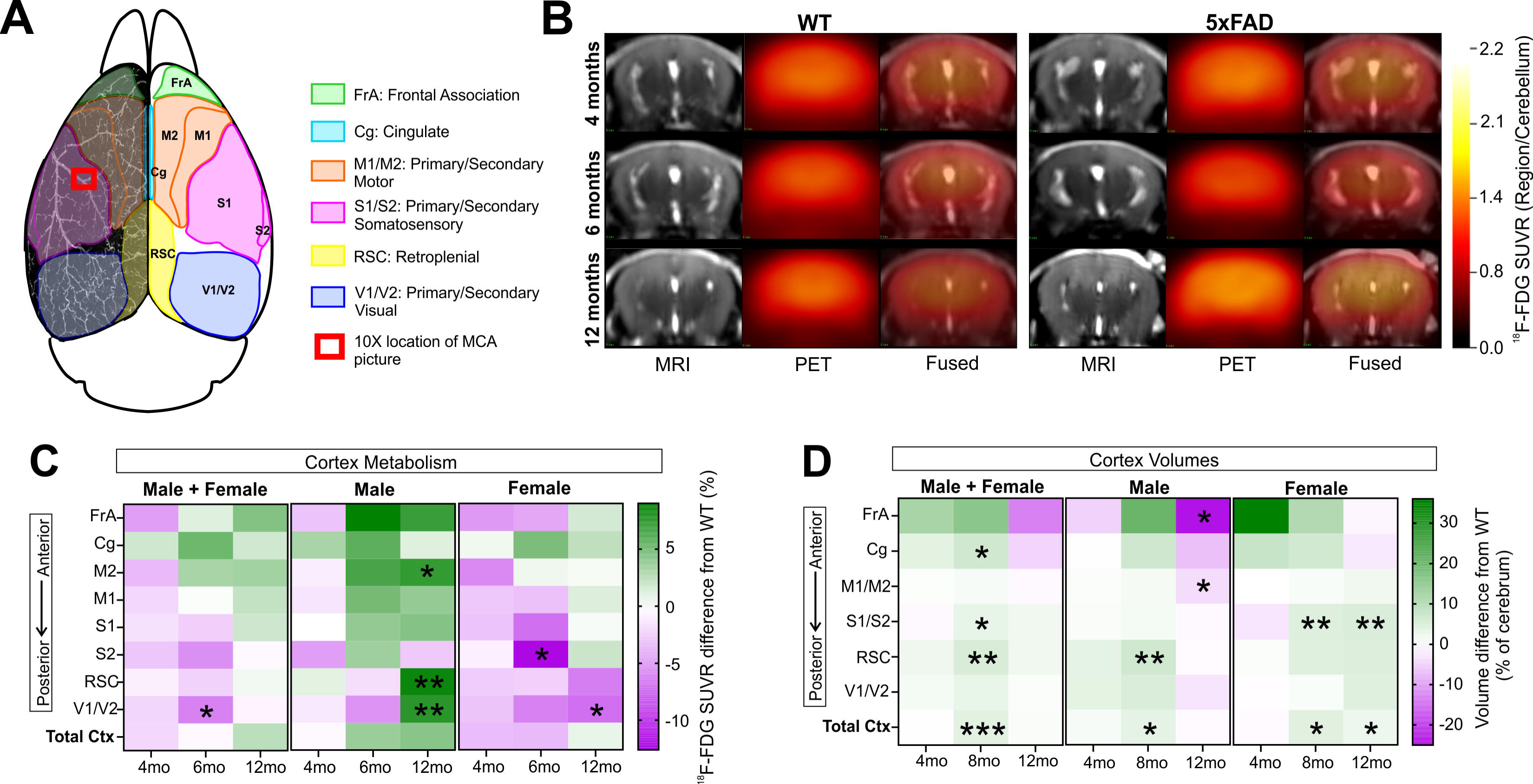
18F-FDG PET cortical metabolism. Schematic of the cortical surface showing the regions analyzed which match the vascular topology measures **(A)**. Representative 18F-FDG PET/MRI images from 6 randomly selected females WT and 5xFAD, at 4, 6, and 12 months of age (Bregma level -1.94mm). PET images (center) are presented as standardized uptake value ratio (SUVR, to cerebellum). PET/MRI fused images are shown on the right **(B)**. Cortical metabolic changes in 5xFAD compared to WT mice **(C)** where male 5xFAD mice exhibited a temporal progression of increased metabolism that was not apparent in female cortical regions. In 5xFAD male mice at 12 months of age, the M2, RSC and V1/V2 cortical areas had significantly increased 18F-FDG uptake in contrast to female mice which exhibited decreased uptake, suggesting sex differences, (see Supplementary Figure 4). Cortical region volumes were increased in both male and female 5xFAD mice compared to WT **(D)**. T-tests compared WT and 5xFAD mice with *=p<0.05, **=p<0.01, ***=p<0.001. **(A)** is modified from (Kirkcaldie, 2012). FrA: frontal association, Cg: cingulate, M1/M2: primary/secondary motor area, S1/S2: primary/secondary sensorimotor area, RSC: retrosplenial dysgranular cortex, V1/V2: primary/secondary visual area.

We next assessed cortical region volume changes in 5xFAD and WT mice to determine if there was an overlap between brain volume and ^18^F-FDG uptake (Figure 9D). In both males and females there was a significant increase in total cortical volumes at 8 months of age of the 5xFAD mice compared to WT, notably with larger posterior regions compared to anterior. At 12 months of age, 5xFAD male mice had significant decreased volume regions in the frontal association (FrA) and primary and secondary motor cortices (t-test, frontal association area: * p=0.022, motor area: * p=0.024, cingulate cortex: p=0.08, Figure 9D). In 12-month-old 5xFAD females, the primary and secondary somatosensory cortex volume was increased compared to WT (** p=0.006), as well as the total cortex volume (* p=0.037). In summary, the modest posterior cortical volume increases at 8 months may drive the increased metabolic demand observed at 12 months of age in the 5xFAD mice, specifically in males.

## 4 Discussion

The neurovascular compartment is thought to play an important role in the onset, evolution, and pathogenesis of AD. In male and female 5xFAD mice across their lifespan, we assessed vascular network characteristics, cortical region volumes and metabolic alterations. Broadly we report the following, as summarized in Table 3: 1) cerebrum volumes were increased with age in WT and 5xFAD mice in both males and females, but significant differences in genotypes were observed only in 8-month-old females where WT cerebrum volumes were significantly larger compared to 5xFAD (Figure 1); 2) Axial cortical surface vessel characteristics (vessel density, junction density, average and total vessel length) were increased with age in both genotypes (Figure 2), but this was driven by males (Figure 3); 3) MCA vessel characteristics showed age and genotype effects, again predominately in males (Figures 5, 6); 4) BBB disruption increased with age in both males and females with a significant genotype effect at 8 months, where half of the 5xFAD mice presented with altered BBB, but WT did not (Figure 7); 5) Vessel collaterals in the watershed were decreased with age in both genotypes independent of sex. Vessel diameters of collaterals reported differing patterns with age and by genotype across males and females (Figure 8); and 6) Glucose metabolism in cortical regions exhibited increased utilization with age in 5xFAD males whereas female mice had reduced ^18^F-FDG uptake, especially at 12 months of age (Figure 9). In summary, vascular alterations and glucose metabolism are dynamically altered with age and across sex in the 5xFAD mouse model of AD compared to age-matched WT mice.

**Table 3:**
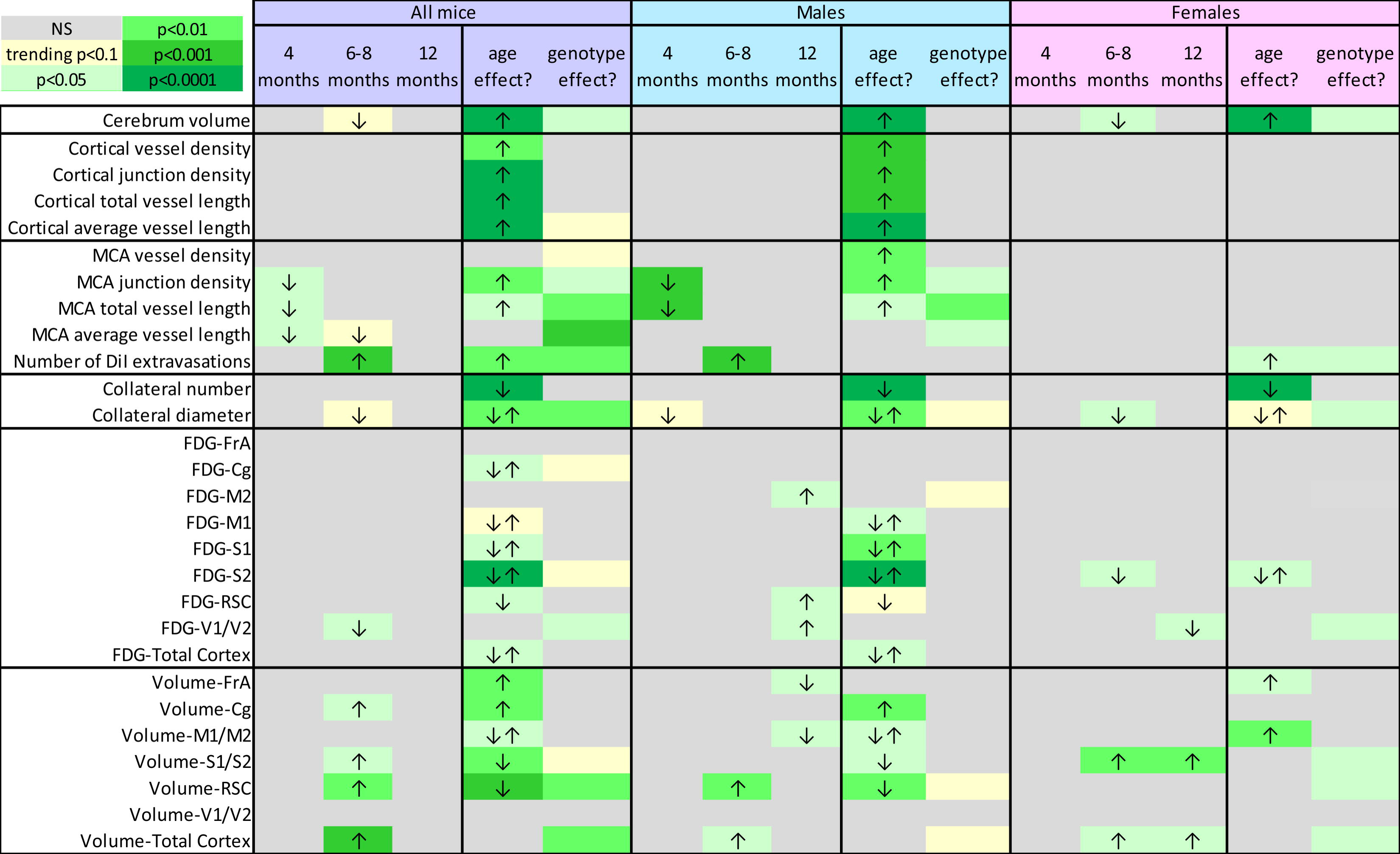
Summary of the results as change in 5xFAD mice compared to WT.

In our recent review we noted that few studies assessed brain volume changes in male and/or female 5xFAD mice longitudinally (Jullienne et al., 2022b). 5xFAD and WT mice at 2-, 4- and 6-month-old, reported no differences in total forebrain, cerebral cortex or frontal cortex volumes (Girard et al., 2013; Girard et al., 2014). These studies did not directly test for sex differences and did not test mice older than 6 months. Although different from human physiology, the increase of cerebrum volumes with age in our 5xFAD study is consistent with other mouse MRI studies which revealed enlarged brain volumes with aging: between 6 and 14 months of age in WT (C57BL/6) and APPxPS1 males (Maheswaran et al., 2009), and between 4 and 24 months in WT and PS2APP females (von Kienlin et al., 2005). To our knowledge, no study has examined sex differences in the increased brain volume during aging. In 8-month-old WT mice, we found an increased brain volume in females compared to males that plateaued at 12 months of age with no significant differences late in life.

It is remarkable that very few studies have examined vascular changes in the 5xFAD mouse (Szu and Obenaus, 2021). Two studies used glucose transporter-1 (GLUT1) immunostaining and observed decreased number of vessels in cortex and hippocampus of 9-month-old female 5xFAD mice compared to WT (Ahn et al., 2018), and decreased capillary lengths in the cortex of 8-month-old 5xFAD females (Kook et al., 2012). It is important to note that our results show vascular differences on the axial cortical surface (Figures 2, 3) or from the M3 portion of the MCA (Figures 5, 6). Our large-scale hemispheric analyses of the cortical surface did not find any differences between WT and 5xFAD mice for any of the vascular metrics nor between sexes. One reason for no genotype or sex differences could be that we sampled the entire axial surface (∼90 mm^2^) and thus regional decreases in specific cortices would be potentially minimized in the 5xFAD mice (Figure 2D). In support of this possibility, the MCA analysis observed significant decreases in junction density and vessel length at 4 months in male 5xFAD mice compared to WT, while female mice showed no differences across age.

A recent review of clinical and preclinical studies found that most results to date report no changes or a modest decrease in vessel density in the presence of AD pathology (Fisher et al., 2022). Broadly, most studies focused on either male or female mice without explicitly testing for sex differences. In regards to the effect of aging alone on vascular density, many studies showed a decrease with age in both humans (Bell and Ball, 1981; Brown and Thore, 2011) and rodents (Sonntag et al., 1997; Wang et al., 2022) which differs from our results with increasing vascular density with age in both WT and 5xFAD mice. However, Giuliani and colleagues reported that cortical microvessel density, as evaluated by laminin immunostaining on coronal slices, was higher in 5-month-old Tg2576 mice, compared with age-matched WT, but was lower at 27 months in Tg2576 mice (Giuliani et al., 2019). We also reported an overall increase in vascular density with age in the 3xTg-AD mouse model using vessel painting (Jullienne et al., 2022a). Thus, across the cortex of WT and 5xFAD mice there is a generalized increase in vascular features, which appears to be independent of genotype when both sexes are combined. Fisher and colleagues also state in their review the discrepant data on vessel density in AD models that likely reflect different methods of analysis, metrics assessed, regions of interest and other factors such as age and sex (Fisher et al., 2022). For example, in non-transgenic adult mice, vessel density has been shown to be higher in primary sensorimotor cortex, where the need for energy support is more important, compared to cortical association areas (Wu et al., 2022). Here, the cortical surface vessel density results are consistent with our previous results in C57BL/6J mice where we reported a density of about 20% in 6-7-month-old males and females (Jullienne et al., 2018). It is noteworthy that our vessel painting method stains the entire cerebral vasculature and our axial images capture vessel features across a 1 mm deep cortical slab while vessel density reported from immunohistochemistry typically report from 10-30μm thick cortical slices.

To our knowledge, no studies have assessed cerebrovascular complexity using fractal geometry in models of AD. The fractal nature of pial vasculature has been reported in cats (Hermán et al., 2001) and we have used it to assess brain vasculature in rodent models of traumatic brain injury (Obenaus et al., 2017; Jullienne et al., 2018). Our results show a significant increased complexity of the vascular network in 8-month-old 5xFAD females compared to WT. Fractal analysis could be used as a potential biomarker, as suggested in a recent human cerebral small vessel disease (CSVD) study, which found that the complexity of the circle of Willis was lower in asymptomatic CSVD patients compared to healthy controls (Aminuddin et al., 2022). Additional studies are needed to determine how vascular complexity is modulated by AD, aging, and sex, but fractal analysis of the cerebrovasculature based on non-invasive MRI angiography could assist in detecting putative vascular dysfunction.

A hallmark of AD pathology is an altered BBB due to pathogenesis and resulting inflammation (Sweeney et al., 2019). BBB disruption has been quantified in 5xFAD mice using several methods. An Evans blue assay used by Ries and colleagues showed that BBB permeability was increased in 3-month-old 5xFAD male mice compared to WT, but at 6 months there were no significant differences (Ries et al., 2021). Alternatively, fibrinogen immunostaining in the cortex and hippocampus of 6-month-old females showed increased BBB permeability in 5xFAD compared to WT mice (Mota et al., 2022). Expression of Zonula Occludens-1 (ZO-1), a tight junction protein, revealed a 60% down-regulation in 9-month-old 5xFAD females compared to age-matched controls (Ahn et al., 2018). In female 5xFAD mice, FITC-albumin leakage representative of microvascular damage, was absent at 2 months of age, first appearing at 4 months and becoming more prevalent by 9 and 12 months (Giannoni et al., 2016). In the hAPPJ20 mouse line, albumin extravasation started at 3 months and increased with age. Moreover, the tight junction protein claudin-5 was down-regulated compared to WT littermates, starting at 2 months in the hippocampus and 3 months in the cortex (Shibly et al., 2022). In summary, there is considerable evidence that BBB elements are modified with increasing age in most AD mouse models that lead to vascular leakage.

We also observed a progressive increased BBB leakage with age in the 5xFAD mice based on vascular DiI extravasation, where male 5xFAD mice exhibited markedly and sustained leakage while female mice had a significant decrease in the number of leaks between 8 and 12 months of age. A question remains if these regions of DiI extravasation are due to permeability changes in the BBB or if they represent microbleeds. There is literature supporting the occurrence of both, BBB disruptions (Zipser et al., 2007; Sweeney et al., 2018) and microbleeds (Akoudad et al., 2016; Graff-Radford et al., 2020) in human AD. Similar findings have reported BBB disruptions (Montagne et al., 2017) and microbleeds (Reuter et al., 2016; Cacciottolo et al., 2021) in mouse of models of AD. Microbleeds in AD have been shown to increase with age both in clinical (Graff-Radford et al., 2020) and in preclinical studies in APP23 (Reuter et al., 2016) and in 5xFAD mice (Cacciottolo et al., 2021). Microbleeds were present at 2 and 4 months of age. At 6 months, 5xFAD females had 30-50% more microbleeds than males, and APOE4 5xFAD females had 25% more microbleeds than APOE3 5xFAD females (Cacciottolo et al., 2021). The mechanisms underlying this disruption in the BBB with increasing age in AD mouse models warrant future studies.

The watershed zone, where the middle, anterior and posterior cerebral arteries intersect, is a known region of vulnerability, particularly in injuries such as stroke (D’Amore and Paciaroni, 2012). Given the susceptibility of this region, we sought to characterize the angioarchitecture of these collateral vessels. A key finding in our 5xFAD mice was that there were no differences in sex, but there was a precipitous decline in the number of collaterals at 12 months of age compared to 4 and 8 months. In normal C57BL/6J mice, there was a decrease in pial collateral vessels extent (number and diameter) with aging (Faber et al., 2011). Collateral rarefaction (decreased collateral density) was assessed in different mouse models of AD (single: APPSwDI, double: APP695, PSEN-1 and triple transgenic: APPSw, PSEN-1, MAPT). WT and single transgenic did not exhibit collateral rarefaction whereas double and triple transgenic mice sustained rarefaction at 8 months of age with no progressive increases at 18 months of age (Zhang et al., 2019). These findings are consistent with our results where we did not observe a progressive increase in rarefaction in the 5xFAD mice when compared to WT. However, the average vessel diameter in 5xFAD females at 8 months of age was significantly decreased compared to WT. Globally, our results suggest that aging has a larger impact on collateral rarefaction than genotype.

Considerable studies in human AD have reported decreased glucose metabolism (^18^F-FDG uptake), in cortical regions such as the cingulate, precuneus and frontal cortices (Minoshima et al., 2021). Other brain regions exhibit relatively well-preserved glucose uptake, albeit individual studies vary in their findings. ^18^F-FDG uptake is considered a reliable marker for AD onset particularly for confirmation of onset of mild cognitive decline. Preclinical studies in mouse models of AD have also been useful in corroborating human AD findings, as recently reviewed (Bouter and Bouter, 2019). However, the review notes contradictory findings that are dependent upon the AD mouse model being investigated. In 5xFAD mice, increased brain metabolism was reported at 11 months of age (Rojas et al., 2013) whilst decreased brain metabolism was found at 13 months of age (Macdonald et al., 2014). At 7 and 12 months of age, 5xFAD male mice had reduced ^18^F-FDG whole brain uptake. In male 5xFAD mice at 12 months, virtually every region tested exhibited metabolic reductions (Franke et al., 2020) and this was also reported in female 5xFAD mice (Bouter et al., 2021). Alternatively, Choi and colleagues reported increased hippocampal ^18^F-FDG uptake compared to WT (Choi et al., 2021).

Our own studies herein found that there was increased metabolic uptake in selected cortical regions in males and modest reductions in ^18^F-FDG uptake in females, consistent with the notion of regional sensitivity. These increases in metabolic activity coincided with a modified angioarchitecture that we described and as we have previously reviewed (Szu and Obenaus, 2021). Cerebral hypoperfusion has been reported in the human AD literature, using [^99m^Tc]HMPAO PET imaging for cerebral perfusion and no overt decrements were noted in relative cerebral blood flow in 5xFAD mice (DeBay et al., 2022). Others have reported no changes in CBF at 12 months (hypoperfusion was present at 7 months) but modestly increased cerebral blood volume (CBV) associated with no changes in glucose metabolism (Tataryn et al., 2021). Clearly there is divergent literature on the consequences of vascular alterations in the 5xFAD mouse model. An important caveat is that there is no uniform method of analyses across many of the PET studies with some using glucose concentrations to correct for uptake and others using regions relative to another region (i.e. cerebellum). It is highly probable that some of the variance in these findings are due to methodological approaches and some consensus in the neuroimaging field would potentially reduce these somewhat disparate findings, as has been recently proposed for MRI (Jelescu et al., 2022; Schilling et al., 2022).

There are several limitations in the current study. One limitation of the vessel painting approach is that animals are sacrificed at each time point thereby providing a cross-sectional view while a longitudinal study could report the vascular alterations across each individual subject. Several non-invasive methods, such as angiography or laser doppler studies could provide these in-vivo assessments albeit at much lower resolution than microscopy of the vessels. Further, as noted above we only assessed the vessels on the cortical surface but examination of deeper structures, such as the hippocampus and temporal lobes, could provide significant information on vascularity that may underlie the cognitive decline reported in the 5xFAD mouse model of AD. An additional strength would be to undertake the metabolic and the vessel phenotyping studies in the same mouse and there are plans to do so in future studies.

In summary, we describe lifespan modifications of the cortical vasculature and glucose metabolism of the 5xFAD mouse model of AD spanning 4 and 12 months of age and across sex (Table 3). We found that increasing age resulted in an increase in cerebrum volumes and in vessel characteristics of the cortical surface, which were not modulated by genotype but exhibited sex differences. MCA vessel characteristics were influenced by age and genotype and were driven by males. BBB disruption was also increased with age and increasing AD pathology worsened leakage. Collateral numbers decreased with age independent of genotype. Finally, glucose utilization in cortical regions was differentially altered by sex in 5xFAD mice. These data confirm the involvement of cerebral vasculature in AD and most importantly highlight the need to report and consider age and sex of the subjects used in studies.

## 7 Author Contributions

AJ, JIS, PRT and AO contributed to conception and design of the study. AJ, JIS, RQ, MT, TN, BPN, AAB, KE, SCP, PRT, and SCP collected and compiled the data. AJ, JIS, RQ, BPN, SCP, PRT and AO analyzed the data. AJ, JIS, RQ, BPN, SCP, PRT and AO did statistical analyses. AJ and AO made figures. AJ, AO and PRT wrote the first draft of the manuscript. All authors contributed to manuscript revision, read, and approved the submitted version.

## 9 Acknowledgments

The authors acknowledge all members of the UCI and IU MODEL-AD consortia for their discussions related to the data reported here. The authors also acknowledge the Stem Cell Research Center Core and their staff at UCI for training and use of their equipment (FV3000 microscope; for the images in Figure 6), the University of Calgary, Experimental imaging centre, and Loma Linda University Center for Imaging Research for MRI acquisition.

## 10 Data Availability Statement

The raw data supporting the conclusions of this article will be made available by the authors, without undue reservation.

## 11 Tables

Table 3 sent as tiff file

