## Supplementary material for "Cortical cerebrovascular and metabolic perturbations in the 5xFAD mouse model of Alzheimer’s disease": Suppl. Fig 1 to 4

### Supplementary Figures

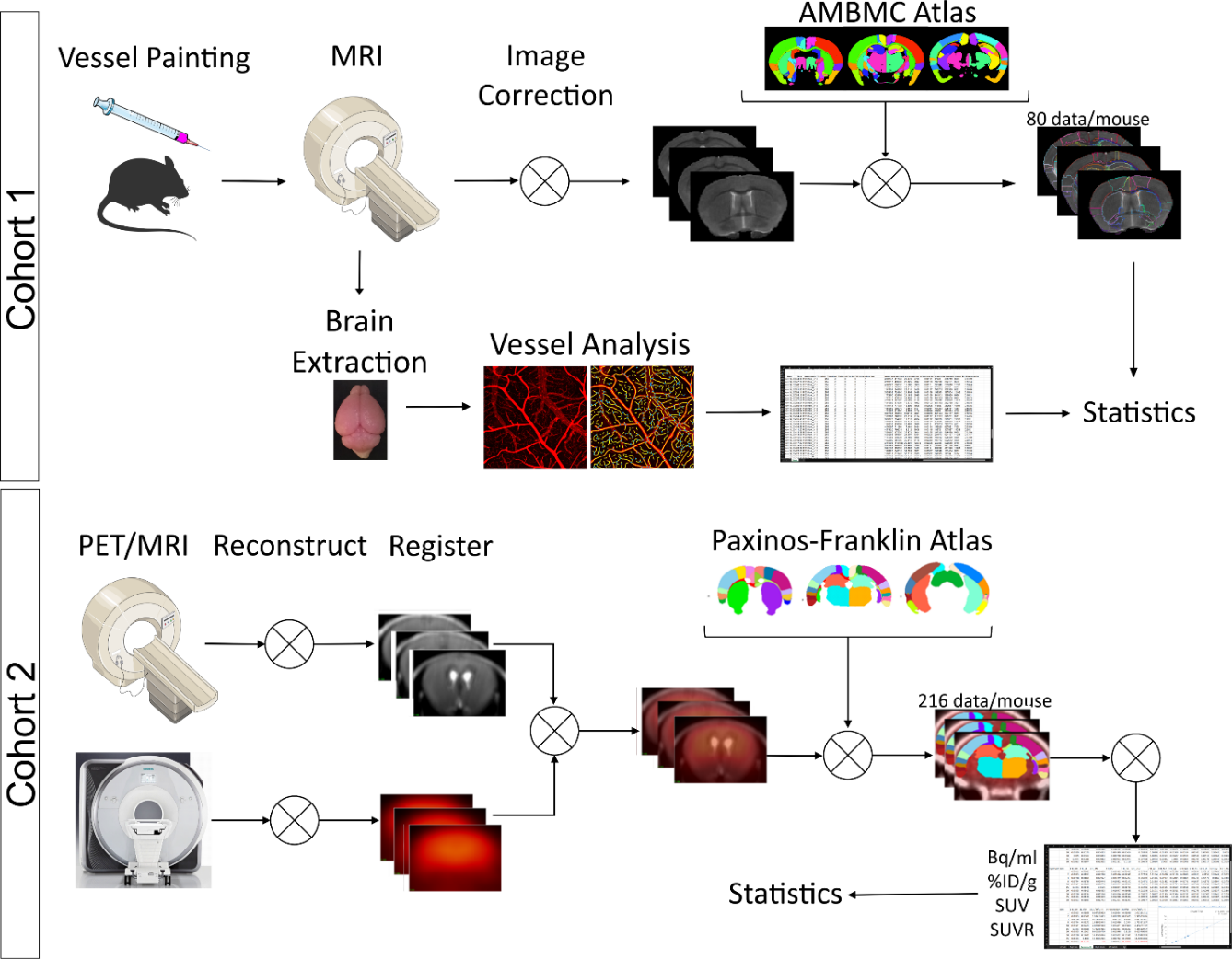

**Supplementary Figure 1:** Experimental design. Two cohorts of age- and sex-matched 5xFAD and WT mice were used for independent experiments. Cohort 1 (4-, 8-, and 12-month-old mice) underwent vessel painting followed by high-resolution ex vivo MRI for vessel network and regional brain volumes analysis. Cohort 2 (4-, 6-, and 12-month-old mice) went through ^18^F-FDG-PET/MRI for regional brain metabolic analyses.

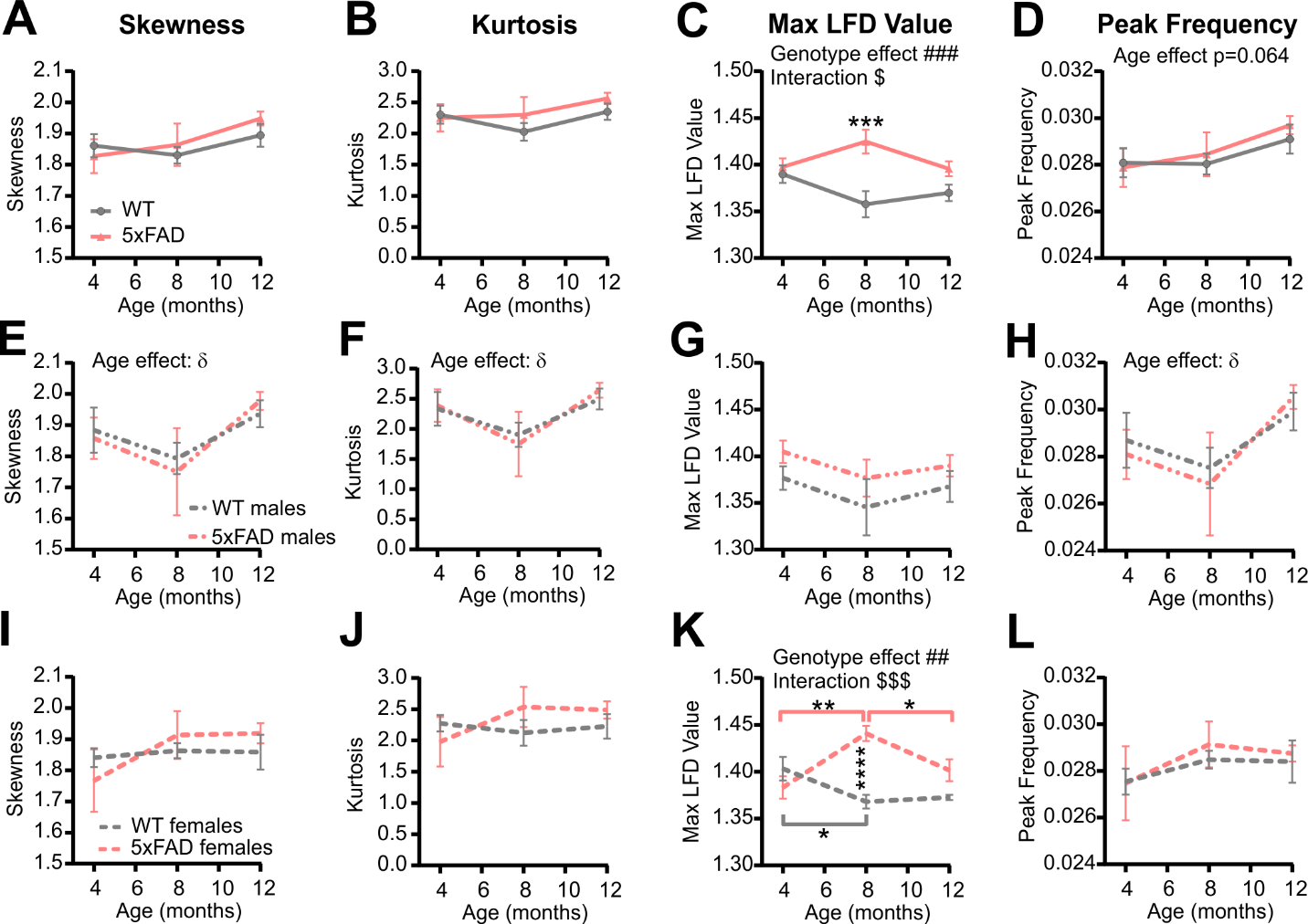

**Supplementary Figure 2:** Metrics associated with the local fractal dimension (LFD) histograms (see Figure 4). Skewness, kurtosis, maximum LFD value, and peak frequency values are shown for males and females combined **(A-D)**, for males only **(E-H)**, and females only **(I-L)**. δ shows a significant effect of age (two-way ANOVA, δ=p<0.05); # shows a significant effect of the genotype (two-way ANOVA, ###=p<0.001); for multiple comparisons across ages and between genotypes (Sidak’s test): **=p<0.01, ***=p<0.001, ****=p<0.0001.

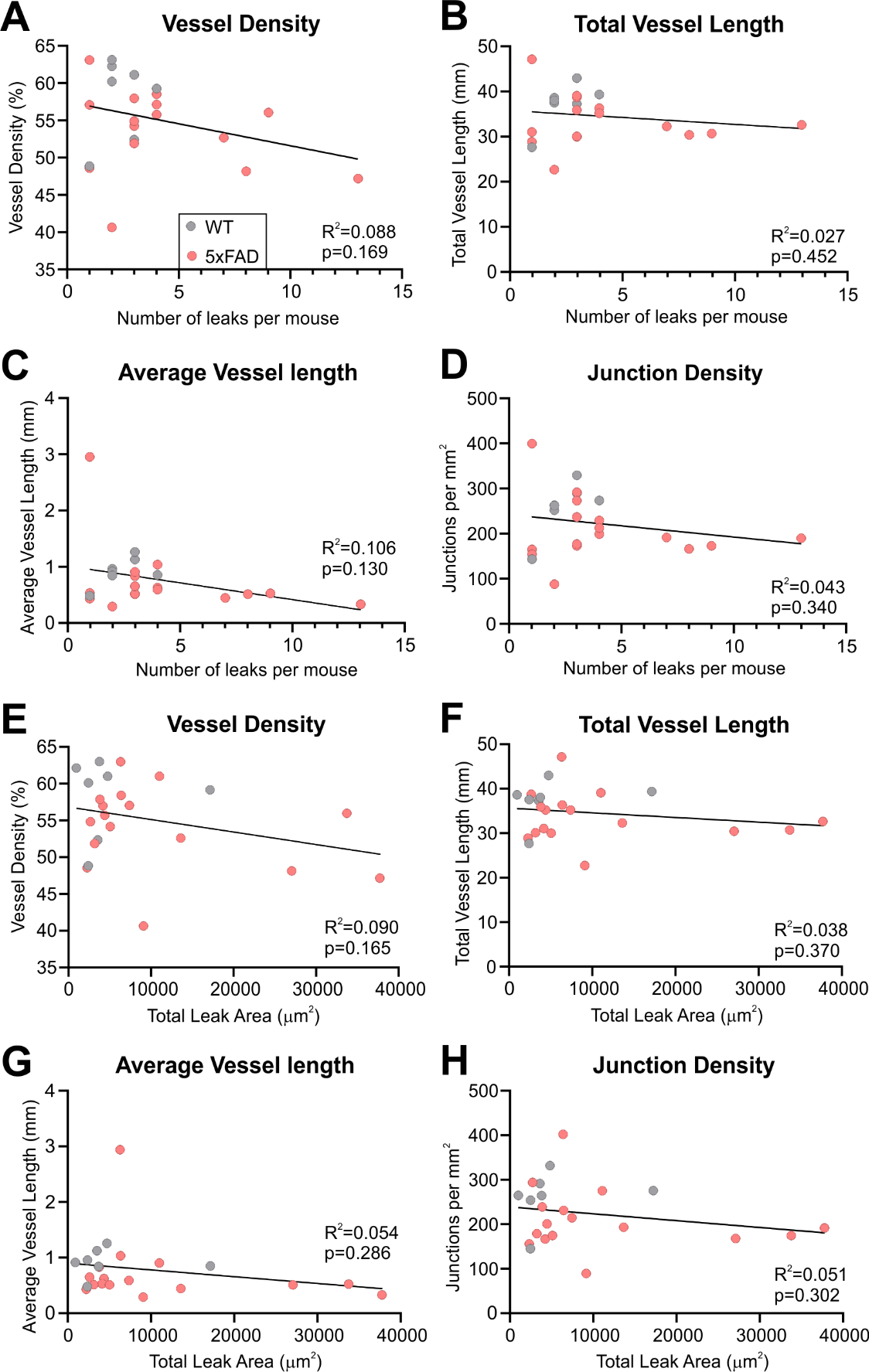

**Supplementary Figure 3:** Correlation plots between classical vessel metrics and number of DiI leaks per mouse (**A-D**) or total leak area per mouse (**E-G**). No correlations were significant. All data were assessed using linear regression modeling and goodness of fit function in the GraphPad software (version 9.5.1).

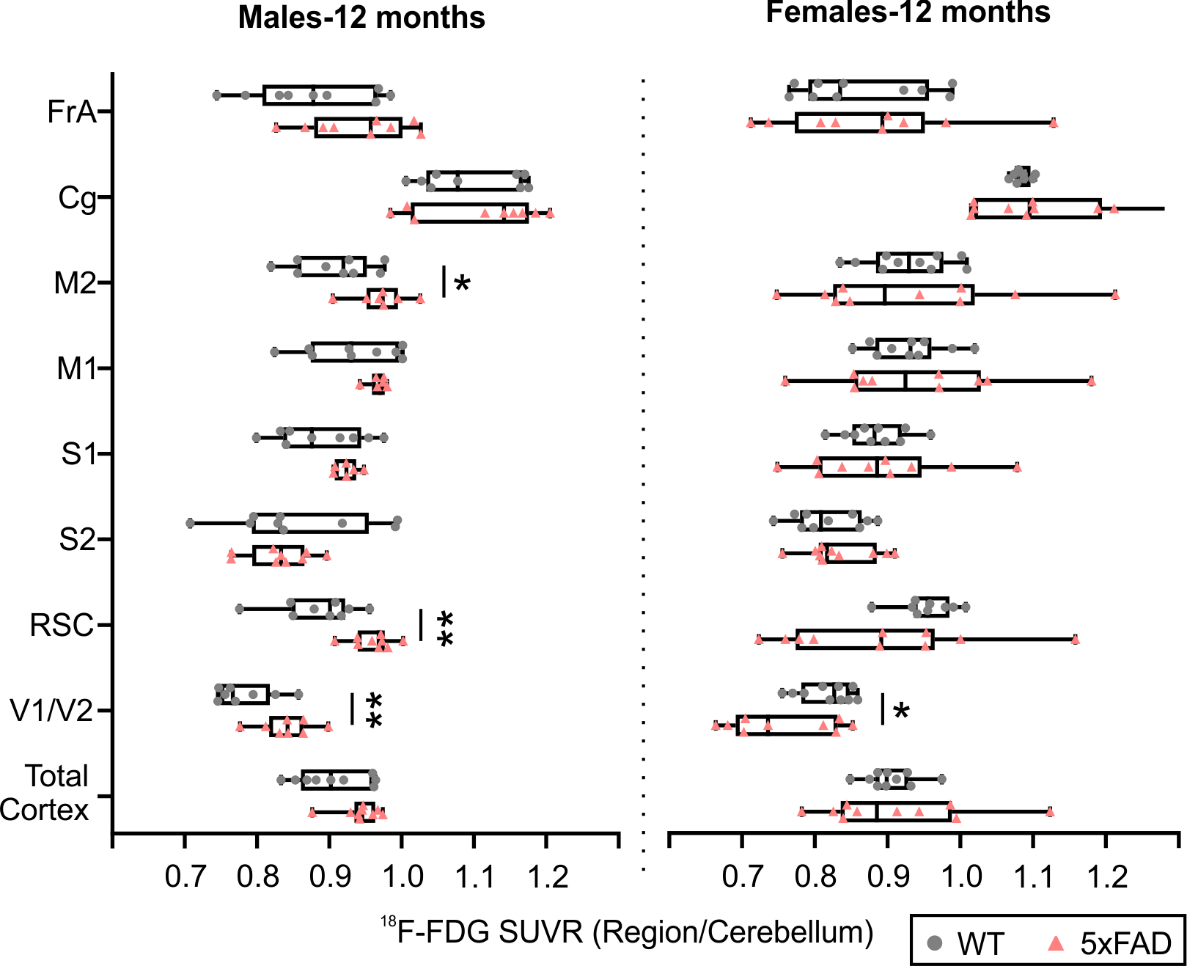

**Supplementary Figure 4:** ^18^F-FDG PET cortical metabolism. In 5xFAD male mice at 12 months of age, the M2, RSC and V1/V2 cortical areas had significantly increased 18F-FDG uptake in contrast to female mice which exhibited decreased uptake, suggesting sex differences. T-tests compared WT and 5xFAD mice with *=p<0.05, **=p<0.01, ***=p<0.001. FrA: frontal association, Cg: cingulate, M1/M2: primary/secondary motor area, S1/S2: primary/secondary sensorimotor area, RSC: retrosplenial dysgranular cortex, V1/V2: primary/secondary visual area.
